# A Non-genetic Mechanism for Chemoresistance in Lung Cancer: The Role of Integrin β4/Paxillin Axis

**DOI:** 10.1101/781807

**Authors:** Atish Mohanty, Arin Nam, Alex Pozhitkov, Supriyo Bhattacharya, Lu Yang, Anusha Nathan, Xiwei Wu, Saumya Srivastava, Isa Mambetsariev, Michael Nelson, Rajendra Pangeni, Dan Raz, Yihong Chen, Yanan He, John Orban, A. R. Subbalakshmi, Linlin Guo, Mohd W. Nasser, Marianna Koczywas, Surinder K. Batra, Mohit Kumar Jolly, Prakash Kulkarni, Erminia Massarelli, Ravi Salgia

## Abstract

Tumor heterogeneity and cisplatin resistance are a major cause of tumor relapse and poor survival. Here we show that in lung adenocarcinoma (LUAD), paxillin (PXN) and integrin beta 4 (ITGB4) are associated with tumor progression, and cisplatin resistance. Silencing PXN and ITGB4 render cisplatin tolerant cells sensitive, and immunologically neutralizing ITGB4 improves sensitivity. The N-terminal half of PXN is intrinsically disordered and interacts with ITGB4 to regulate expression of USP1 and VDAC1 which are required for maintaining genomic stability and mitochondrial function in LUAD. By virtual screening an FDA-approved compound library, we identified compounds that interact with PXN *in silico* and attenuate cisplatin resistance in LUAD cells. RNAseq analysis identified a double negative feedback loop between ITGB4 and microRNA miR-1-3p, suggesting that bistability could lead to stochastic switching between cisplatin-sensitive and resistant states in these cells. The data highlight an alternate, non-genetic, mechanism underlying chemoresistance in lung cancer.

## Introduction

Lung cancer is the most frequently diagnosed cancer and a leading cause of cancer-related deaths worldwide (Bray et al, 2018). Approximately 85% of patients have a group of histological subtypes collectively known as non-small cell lung cancer (NSCLC), of which lung adenocarcinoma (LUAD) and lung squamous cell carcinoma are the most common subtypes, followed by squamous cell carcinoma and less so, large-cell carcinoma (Salgia, 2016; Herbst et al, 2018). LUAD accounts for ∼40% of all lung cancers. These histologies possess different clinical characteristics, and there are potential differences in response to cytotoxic chemotherapies. Approximately 40–50% of patients with NSCLC will be diagnosed with advanced or metastatic disease and are not candidates for curative therapy. Recent advances have transformed lung cancer care with a percentage of patients receiving first-line tyrosine kinase inhibitors based on their genomic-informed markers (Tan et al, 2017). However, immunotherapy alone or in combination with platinum-based chemotherapy is now the recommended first-line treatment option for the remainder of these patients.

Cisplatin [Cis-diamminedichloroplatinum(II)] is a widely prescribed platinum-based compound that exerts clinical activity against a wide spectrum of solid neoplasms, including testicular, bladder, ovarian, colorectal, lung, and head and neck cancers (Galluzzi et al, 2012; Galluzzi et al, 2014). Cisplatin treatment is generally associated with high rates of clinical responses. However, in the vast majority of cases, malignant cells exposed to cisplatin activate a multipronged adaptive response that renders them less susceptible to the anti-proliferative and cytotoxic effects of the drug, and eventually resume proliferation. Thus, a large number of cisplatin-treated patients experience therapeutic failure and tumor recurrence. However, the exact mechanism(s) underlying the emergence of drug resistance remains poorly understood and a bewildering plethora of targets have been implicated in different cancer types. For example, in NSCLC alone, dysregulation of genes involved in cell cycle arrest and apoptosis namely, mouse double minute 2 homolog (MDM2), xeroderma pigmentosum complementation group C, stress inducible protein and p21 (Sarin et al, 2017), cytoplasmic RAP1 that alters NF-κB signaling, upregulation of antiapoptotic factor BCL-2 (Xiao et al, 2017), enhanced Stat3 and Akt phosphorylation, high expression of survivin (Hu et al, 2016), hypoxia factor HIF-1α and mutant p53 (Deben et al, 2018), have been implicated in cisplatin resistance. Furthermore, although changes in administration schedules, choice of methods, and frequency of toxicity monitoring have all contributed to incremental improvements, chemoresistance limits the clinical utility of cisplatin (Fennell et al, 2016). Therefore, there is a dire need for a deeper understanding of chemoresistance and the identification of prognostic and predictive markers to discern responders from non-responders.

Here, we interrogate the role of the focal adhesion (FA) complex in cisplatin resistance. The FA complex is a large macromolecular assembly through which mechanical force and regulatory signals are transmitted between the extracellular matrix and an interacting cell (Chen et al, 2003). Paxillin (PXN), integrins, and focal adhesion kinase (FAK) are among the major components of this complex. Human PXN is a 68 kDa (591 amino acids) protein (Salgia et al, 1995) and is a recognized contributor to cisplatin resistance in lung cancer (Wu et al, 2014). The N-terminus contains a proline-rich region that anchors SH3-containing proteins and five leucine-rich LD domains (LD1–LD5) with a consensus sequence LD*X*LL*XX*L (Turner, 1998; Turner, 2000; Kanteti et al, 2016). The LD2-LD4 region includes sequences for the recruitment of signaling and structural molecules, such as FAK, vinculin, and Crk. This region has also been reported to interact with integrin α, more specifically, integrin α4 (ITGA4) (Liu et al, 1999; Liu and Ginsburg, 2000). The C-terminal region is also involved in the anchoring of PXN to the plasma membrane and its targeting to FAs. It contains four cysteine-histidine-enriched LIM domains that form zinc fingers, suggesting that PXN could bind DNA and act as a transcription factor. Consistently, PXN is reported to locate to the nucleus which is regulated by phosphorylation (Dong et al, 2009; Ma and Hammes, 2018). In LUAD, expression of PXN is correlated with tumor progression and metastasis (Song et al, 2010; Mackinnon et al, 2011). Further, phosphorylation of PXN activates the ERK pathway, increased Bcl-2 expression, and cisplatin resistance (Wu et al, 2014). Finally, specific PXN mutants, through their interactions with Bcl-2 and dynamin-related protein 1, also regulate cisplatin resistance in human lung cancer cells (Kawada et al, 2013).

Integrins are transmembrane receptors that facilitate cell-extracellular matrix adhesion; they form a critical link between the extracellular matrix and the cell interior by interacting with the FA via PXN. Upon ligand binding, integrins activate signal transduction pathways that mediate cellular signals, such as regulation of the cell cycle, organization of the intracellular cytoskeleton, and movement of new receptors to the cell membrane (Giancotti and Rusolahti, 1999; Maziveyi and Alahari, 2107). Integrins are obligate heterodimers of one α and one β subunit. In mammals, there are 24 α and 9 β subunits (Alberts et al, 2014). Among the various β subunits, β1 is ubiquitously expressed in most cell types and can dimerize with multiple α subunits, forming receptors for various matrixes. On the other hand, integrin β4 (ITGB4) is reported to be quite selective and heterodimerizes only with the α6 subunit and binds to laminin (Mainiero et al, 1997). ITGB4 is also unique because of its >1000 amino acid-long cytoplasmic domain compared to ∼50 amino acid-long domain of other β forms (Su et al, 2008). Interaction of ITGA6/B4 and Shc leads to activation of the RAS-MAPK signaling pathway for cell cycle progression and proliferation (Mainiero et al, 1995). ITGA6/B4 can also activate the PI3 kinase pathway followed by Rac1 to promote tumor invasion (Shaw et al, 1997). In NSCLC, the receptor tyrosine kinase MET interacts with ITGA6/B4 and this interaction is required for HGF-dependent tumor invasion (Trusolino et al, 2001). In addition, the presence of the ITGA6/B4 heterodimer in tumor-derived exosomes facilitates the creation of the microenvironment for lung metastasis (Hoshino et al, 2015). Together, these observations underscore the importance of the FA complex in NSCLC pathophysiology. However, how the interactions of the individual components of the FA complex may contribute to cisplatin resistance remains poorly understood.

In this manuscript, we demonstrate that a significant portion of the N-terminal half of PXN is intrinsically disordered and that the interaction between ITGB4 and PXN is critical for cisplatin resistance. ITGB4 and PXN double knockdown down affects the expression of > 300 genes which are constituents of various pathways required for lung cancer proliferation and survival. USP1 and VDAC1 are two of the top ten genes that are downregulated by the double knockdown and are essential for inducing DNA damage repair induced by cisplatin and for maintaining mitochondrial function, respectively. Via an *in silico* screen designed to identify small molecule compounds that can bind to PXN, we obtained candidate drugs from a library of FDA-approved compounds. Several of these compounds are efficacious in alleviating cisplatin resistance in LUAD cell lines expressing high levels of ITGB4/PXN. Furthermore, using tumor-derived exosomes isolated from patient blood, we identified ITGB4/PXN as potential biomarkers for cisplatin response. Finally, RNAseq analysis identified a double negative feedback loop between ITGB4 and the microRNA miR-1-3p, suggesting that bistability can lead LUAD cells to stochastically switch between cisplatin-resistant and sensitive phenotypes, underscoring a non-genetic mechanism driving the evolution of chemoresistance.

## Results

### Upregulation of both PXN and ITGB4 but not PXN alone correlates with cisplatin resistance

In order to identify the genes upregulated in cisplatin resistance in LUAD cell lines, we performed GSEA analysis on RNAseq data from the Molecular Signatures Database v6.2 (MSigDB), a collection of annotated gene sets (Broad Institute) using the Gene Set Enrichment Analysis (GSEA) software. GSEA is a computational method that determines whether an *a priori* defined set of genes shows statistically significant concordant differences between two biological states. We identified a set of eleven genes that were upregulated in cisplatin-resistant LUAD cells compared to sensitive cells (**Supplementary Table 1**). Four of these eleven genes namely, PXN, ITGB4, ITGA7, and Rac Family Small GTPase 1, are constituents of the FA complex activation, formation, and downstream signaling pathways. Therefore, to verify the correlation of these genes with cisplatin resistance, we randomly selected five LUAD cell lines that harbored wild type (WT) KRAS and five that carried a mutant (MT) version of KRAS, treated them with 10 μM cisplatin for 72 h, and determined cell viability using the CCK-8 viability assay. Among the KRAS WT cell lines, H1437 and H1993 were significantly more resistant while amongst the KRAS MT cell lines, H441 and H2009 were relatively more resistant than the other three cell lines (**Fig. 1A**).

**Figure 1.**
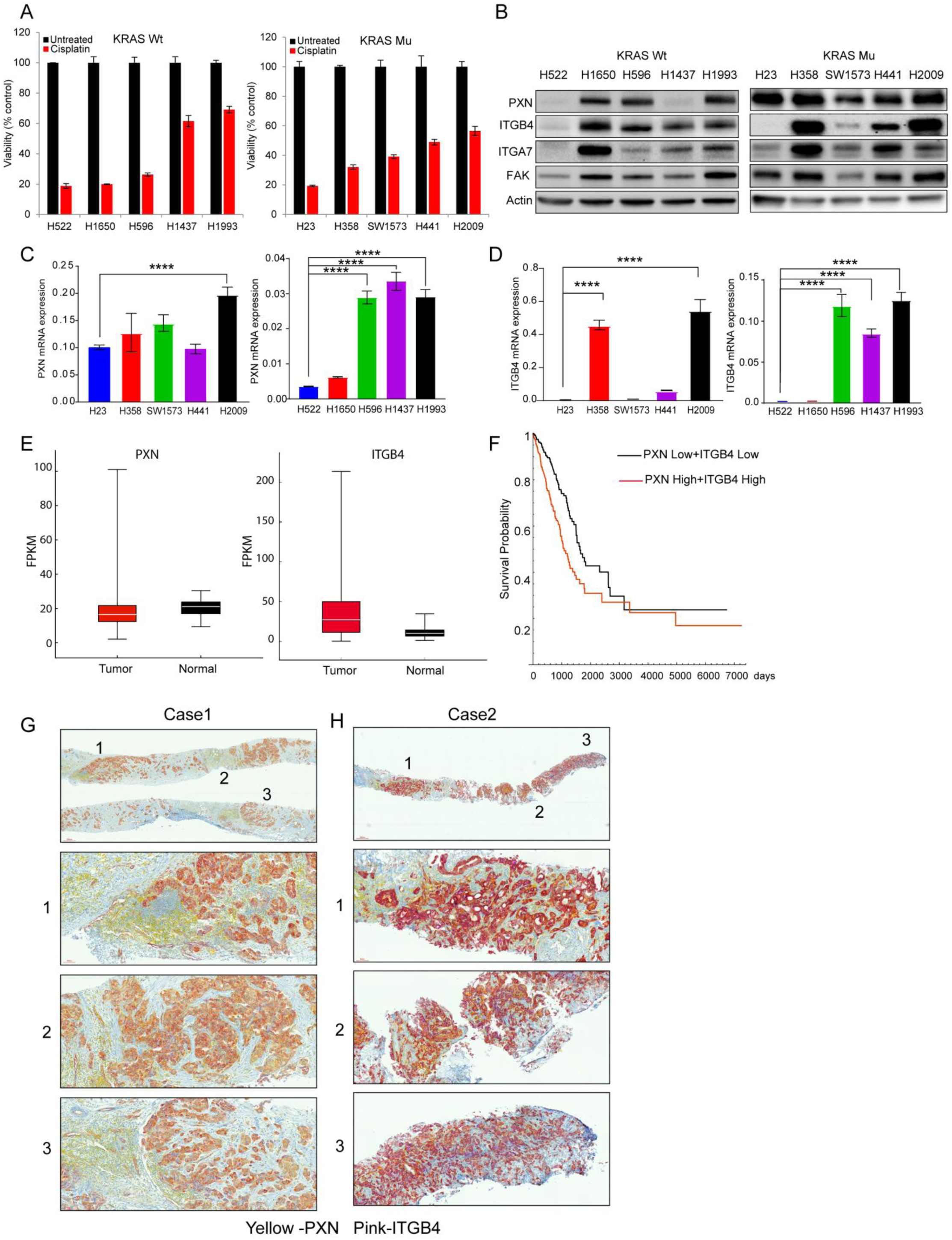
PXN and ITGB4 are upregulated in lung cancer and in cisplatin-resistant LUAD cell lines. **A)** Five LUAD cell lines with wild type (WT) KRAS and 5 cell lines with mutant (MT) KRAS were treated with 10 μM cisplatin for 72 h and cell viability was determined using Cell Counting Kit-8 assay. **B)** Immunoblot showing that the most cisplatin-resistant cell lines (H1993 and H2009) have high expression of both PXN and ITGB4. **C)** and **D)** qPCR results demonstrating high mRNA expression of PXN and ITGB4 in cisplatin-resistant cell lines. **E)** Gene expression profiles of LUAD patients were extracted from TCGA database and expression of PXN and ITGB4 were higher compared to normal tissue. **F)** Kaplan-Meier curves indicate significant survival difference (p=0.000089) between the two groups (PXN low + ITGB4 low and PXN high + ITGB4 high). **G)** and **H)** Immunohistochemistry staining of tissue microarrays showing ubiquitous expression of PXN (yellow) and differential expression of ITGB4 (pink). (**** p<0.0001 One-way ANOVA)

Next, we determined the protein expression of PXN, ITGB4, ITGA7, and FAK in these cell lines by immunoblotting. Rac1 expression was undetectable in these cell lines. However, PXN and FAK were expressed in all the cell lines tested regardless of whether they were cisplatin-sensitive or resistant (**Fig. 1B**). While the expression of ITGB4 and ITGA7 was variable among the cell lines, ITGB4 expression was high in the two cisplatin-resistant cell lines, H2009 and H1993. We also analyzed the mRNA expression patterns encoding these two proteins in these cell lines and confirmed that the mRNA was consistently upregulated for both PXN and ITGB4 in the H2009 and H1993 cell lines (**Fig. 1C & D**).

### Co-expression of PXN & ITGB4 correlates with poorer prognosis in LUAD patients

Since LUAD cell lines overexpressing PXN appeared sensitive to cisplatin treatment when they did not express ITGB4, we hypothesized that there is a correlation between PXN and ITGB4. To test the hypothesis, gene expression profiles of the patients diagnosed with “Lung Adenocarcinoma” were extracted from The Cancer Genome Atlas (TCGA) database and analyzed for expression of ITGB4 and PXN (**Fig. 1E**). The gene-level expression values, FPKM, were analyzed according to the Cox proportional hazards model: Survival ∼ Sex + Stage + Age + PXN * ITGB4. Stage and PXN * ITGB4 showed significant contribution to the model. We separated the patients into two groups by the median value of the PXN*ITGB4 product (**Supplementary Table 2**). The Kaplan-Meier curves showed clear survival difference between the two groups and the log rank test showed significant differences between the two curves (p=0.000089) (**Fig. 1F**). In addition, we performed immunohistochemistry on needle biopsy specimens obtained from 2 de-identified cases from City of Hope. PXN staining was detected using yellow and ITGB4 with pink color. Thus, the coexpression of the two molecules would generate an orange or bright red color based on the expression level. Indeed, we observed significant coexpression of ITGB4 and PXN in various regions of the tumor (**Fig. 1G and H**). The case represented in **Fig. 1G** had weak expression of ITGB4 and coexpression generated an orange color. The second case (**Fig. 1H**) had high expression of ITGB4 and for the same reason it generated a reddish pattern implying coexpression. We further extended the IHC staining to 3 more cases where staining of the whole tumor section was done. We observed variability in the expression of PXN and ITGB4 among cases and within the sample as well. PXN is expressed in most of the region of the tumor but ITGB4 expression is sporadic and the level of ITGB4 expression is also variable among cases. Case 3 had high expression of ITGB4, Case 4 had intermediate expression, and Case 5 had weak expression. PXN expression was seen in the tumors and also in the peripheral region surrounding the tumor tissue that is lymphoid tissue. The normal lung tissue also had weak expression of PXN. The coexpression of PXN and ITGB4 (red or orange color) is limited to some regions of the tumor, emphasizing heterogeneity in their expression patterns (**Supplementary Fig. 1A**). This observation led us to hypothesize cisplatin treatment favors selection of a subclonal population of tumor cells with coexpression of ITGB4 and PXN. Therefore, we interrogated the role of ITGB4 in cisplatin resistance in those cell lines that had high expression of PXN.

### Knocking down ITGB4 attenuates proliferation and migration of cisplatin-resistant cells

We determined the effect of knocking down ITGB4 on cell proliferation using H1993 and H2009 cells. Cells were transiently transfected with a control (scrambled) and ITGB4-specific siRNA, and the effect of silencing its expression on cell proliferation was determined using the IncuCyte Live Cell Imaging System (Essen Bioscience, Ann Arbor, MI). To facilitate live cell analysis, we generated stable cell lines with nuclear expression of the red fluorescence protein (RFP) mKate2 (**Supplementary Fig. 2A**). Knocking down ITGB4 significantly attenuated their proliferation (**Fig. 2A**) and immunoblotting analysis using total cell lysates and qPCR analysis with RNA extracted from transfected cells confirmed a significant decrease in ITGB4 both at the protein and mRNA level (**Fig. 2B**). We also analyzed the doubling time of both cell types. In the case of H2009 cells, we observed that by 50 h post transfection, control (scramble siRNA) cells showed a 4-fold increase in cell count, indicating a doubling time to be approximately 25 h. However, in ITGB4 knocked down H2009 cells, the doubling time increased to 40 h. Similarly, in H1993, knocking down ITGB4 led to only 0.2-fold increase in cell count by 72 h, leading to an estimated doubling time of approximately 120 h compared to 72 h for control cells (**Supplementary Fig. 1B**). We also knocked down integrin A7 in H2009 cell lines to validate its role in cell proliferation but did not observe any significant change (**Supplementary Fig 2D**). Additionally, we also checked for up regulation of expression of other integrin β forms in response to ITGB4 knockdown and found an increase in the expression of ITGB1, ITGB2, and ITGB3 at mRNA level (**Supplementary Fig. 2E**). We further tested the inhibitory effect of ITGB3 inhibitors in the ITGB4 knocked down cells but found no significant difference in their proliferation (**Supplementary Fig. 2F**).

**Figure 2.**
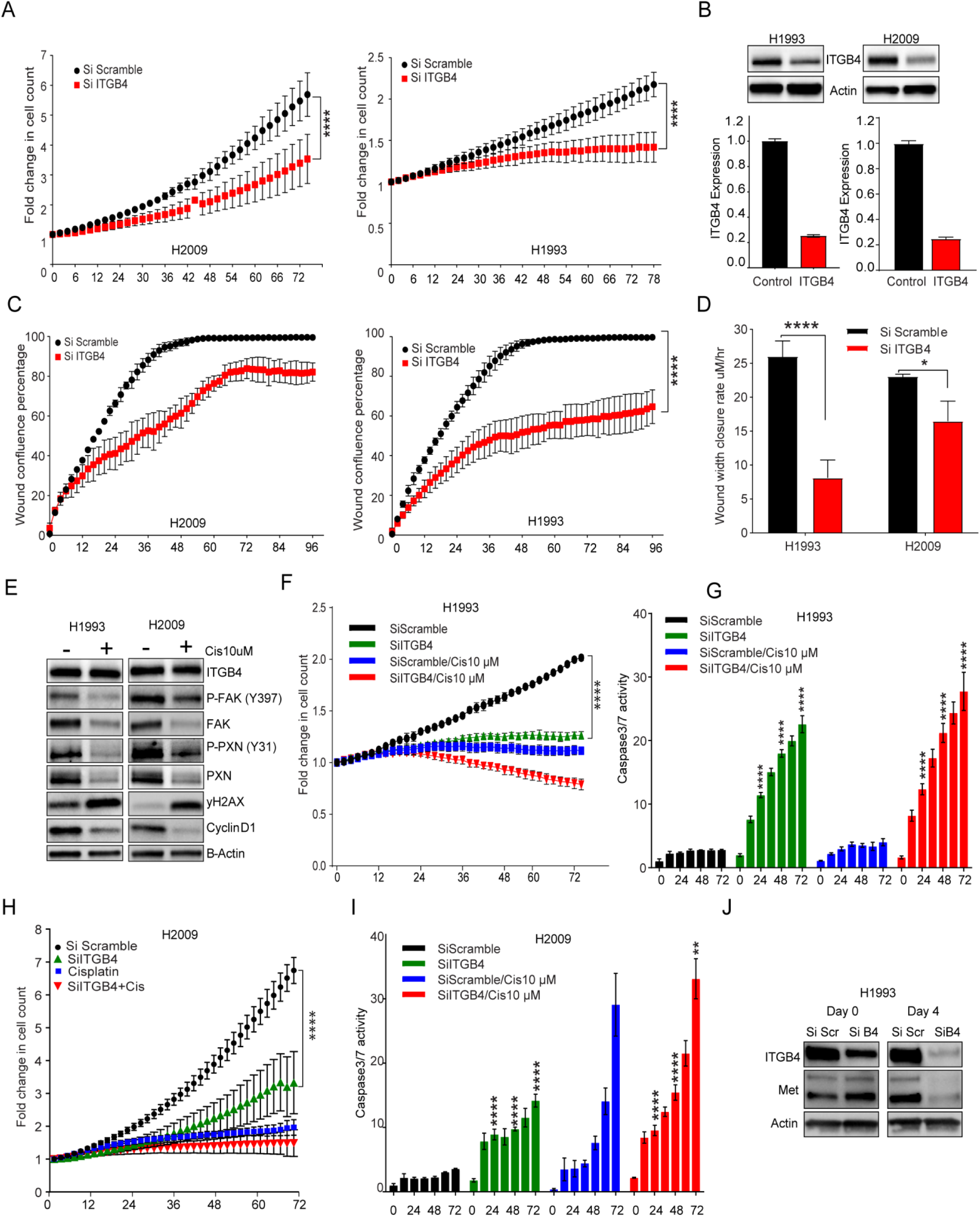
Knocking down ITGB4 attenuates proliferation and migration and increases sensitivity of cisplatin-resistant cells. H2009 and H1993 stable cell lines expressing mKate2 were transfected with control (Si Scramble) or ITGB4-specific (Si ITGB4) siRNA. **A)** ITGB4 knockdown cells (red) had a significantly reduced proliferation rate than control cells (black). (****p<0.0001 Two-way ANOVA) **B)** Immunoblotting and qPCR data confirming the knockdown. **C)** A scratch wound assay demonstrating the effect of knocking down ITGB4. ITGB4 knockdown (red) significantly halted migration and did not close the wound completely after 96 h in both resistant cell lines. (****p<0.0001 Two-way ANOVA) **D)** Rate at which the wound closed was also significantly decreased in ITGB4 knockdown cells (red). (****p<0.0001 and *p<0.0156 Two-way ANOVA). **E)** Cisplatin (10 μM) treatment for 72 h reduced expression in phosphorylated and total FAK and PXN but not ITGB4. **F)** and **G)** ITGB4 knockdown in H1993 cells inhibited proliferation and increased caspase-3/7 activity. Treating ITGB4 knockdown cells with cisplatin had an additive effect to induce caspase activity but drug treatment alone did not have a cytotoxic effect. (****p<0.0001 Two-way ANOVA) **H)** and **I)** ITGB4 knockdown in H2009 cells inhibited proliferation by 72 h but did not induce caspase activity. Cisplatin treatment to ITGB4 knockdown cells for 72 h had an additive effect to induce caspase activity. (****p<0.0001 Two-way ANOVA) **J)** Immunoblot showing MET protein expression in H1993 cells was reduced 4 days after knocking down ITGB4.

To discern the effect of knocking down ITGB4 on cell migration, H1993 and H2009 cells were transfected with the control or ITGB4-specific siRNA in 6-well plates and 12 h post transfection, the cells were trypsinized and reseeded at high density in a 96-well plate. A scratch wound was generated using the WoundMaker from Essen Bioscience and wound healing was observed in real time using the IncuCyte Live Cell Imaging System (**Supplementary Fig. 2C, Supplementary video 1 and 2**). While both cell lines transfected with the scramble siRNA achieved 100% confluency of the wound within 48 h, H1993 and 2009 cells in which ITGB4 was knocked down reached ∼ 60% and ∼ 80% confluency, respectively; neither cell line reached 100% confluency even after 96 h (**Fig. 2C**). We next estimated the rate of wound closure. The rate was determined by dividing initial wound width by the time taken to close the wound. In the case of the H1993 cells, the wound closure rate was ∼ 25 μM/h, and on knocking down ITGB4, the rate significantly dropped to <10 μM/h. Similarly, in the case of the H2009 cells, the wound width closure rate dropped from 22 μM/h to about 18 μM/h (**Fig. 2D**). Together, these results confirmed the significance of ITGB4 in modulating cell proliferation and migration in cisplatin-resistant LUAD cell lines.

### Knocking down ITGB4 increases sensitivity to cisplatin treatment

Next, we determined if inhibition of cell proliferation by knocking down ITGB4 had a cytostatic or cytotoxic effect and whether it was synergistic with cisplatin treatment. To determine apoptosis, we employed the Essen Bioscience Caspase-3/7 Green apoptosis assay in live cells, which uses NucView™488, a DNA intercalating dye, to enable quantification of apoptosis over time. The dye is tagged to DEVD peptide and non-toxic to cells but once the caspase-3/7 is activated, it cleave the dye from the DEVD region. The free dye can intercalate with the DNA leading to the generation of green fluorescence that is then detected in real time and quantified using the IncuCyte System (**Supplementary Fig 3A and Supplementary video 3,4,5 and 6**).

To address this, we treated H1993 and H2009 cells with cisplatin for 3 days and analyzed the changes by immunoblotting. We observed a reduction in the expression of phosphorylated as well as total FAK and PXN, but not in ITGB4 expression (**Fig. 2E**). Next, we transiently transfected H1993 and H2009 cells with ITGB4 siRNA and 24 h post-transfection treated them with 10 μM cisplatin and determined cell proliferation and caspase activity. In H1993, ITGB4 knockdown, or cisplatin treatment, inhibited cell proliferation significantly and the fold change in cell count was ∼1, indicating a cytostatic effect. However, in response to cisplatin in ITGB4 knockdown cells, the fold change dipped to <1, alluding to a cytotoxic effect (**Fig. 2F**). Further, live caspase-3/7 activity analyzed in the same cells showed a 5- to 6-fold increase in activity compared to control by 24 h and by 72 h, caspase activity increased to 20-fold. Addition of cisplatin to the ITGB4 knockdown cells had an additive effect but cisplatin treatment alone did not induce caspase activity, indicating the reduction in cell proliferation by cisplatin was cytostatic but that knocking down ITGB4 induces cytotoxicity (**Fig. 2G**). In KRAS MT H2009 cells, a similar trend was observed but the cell proliferation did not drop <1 and the induction of caspase activity by ITGB4 knockdown was weak. Knocking down ITGB4 induced caspase activity within 24 h of seeding, whereas cisplatin treatment alone induced apoptosis in H2009 cells by 72 h and the combination had an additive effect (**Fig. 2H and I**). ITGB4 knockdown in another KRAS WT cell line H1650 also inhibited cell proliferation and induced cell death. However, cell death was not due to caspase activation but appeared to be due to cell bursting which is related to anoikis (**Supplementary Fig. 3B and Supplementary video 7 and 8**). Together, these data underscore the role of ITGB4 in imparting cisplatin resistance in these two LUAD cell lines irrespective of their KRAS status.

It has been shown that the cytoplasmic domain of ITGB4 acts as a scaffold for various kinases involved in activating the MAPK, PI3k or Akt pathways (Trusolino et al, 2001). MET Proto-Oncogene, Receptor Tyrosine Kinase (MET) is also one of the interacting partners of ITGB4 and is responsible for HGF-dependent phosphorylation and activation of ITGB4 (Trusolino et al, 2001). Of the two cell lines used in the present study, H1993 is known to harbor MET amplification. Thus, we asked if ITGB4 knockdown disrupts MET activity. Indeed, we observed a decrease in MET expression in ITGB4 knocked down cells at the protein level but there was no significant change in the mRNA expression, suggesting that ITGB4 may be involved in MET protein stability in these cells (**Fig. 2J and Supplemental Fig. 3C**). However, in H2009 cells that do not carry MET amplification, no detectable decrease in MET expression was observed (Supplemental Fig. 3D). These data led us to conclude that the cells dependent on MET signaling (H1993) may be more sensitive to ITGB4 knockdown while cells independent of MET signaling (H2009) may be insensitive.

### Treating cells with ITGB4 antibody or knocking down both PXN and ITGB4 has a synergistic effect on attenuating cisplatin resistance

Having demonstrated that silencing ITGB4 expression can increase significant sensitivity to cisplatin, we asked if blocking the ITGB4 extracellular epitope with an antibody would have a similar effect. H1993 cells were first incubated with anti-ITGB4 antibody for 6 h in ultra-low attachment tissue culture plates and then plated on tissue culture-treated plates. As expected, antibody-treated cells were unable to attach to the plate, indicating neutralization of ITGB4 epitopes by the antibody. The cells were allowed to grow for 72 h and proliferation was determined using red fluorescence count. We observed that antibody-treated cells proliferated slower than untreated or control IgG-treated cells. However, ITGB4 antibody treatment induced 5- to 7-fold higher caspase activity than untreated cells (**Fig. 3A and 3B**). Furthermore, immunoblotting performed on whole cell lysates prepared from these cells showed a decrease in ITGB4 and PXN protein levels after 48 h of culturing, which further declined steadily until 72 h (**Fig. 3C**). In contrast, cells incubated with control IgG antibody in ultra-low attachment plates for the same period of time and re-plated in tissue-culture treated plates did not show any change in the ITGB4 protein level, suggesting that decrease in protein levels was due to anti-ITGB4 antibody treatment and not due to any difference in culturing conditions.

**Figure 3.**
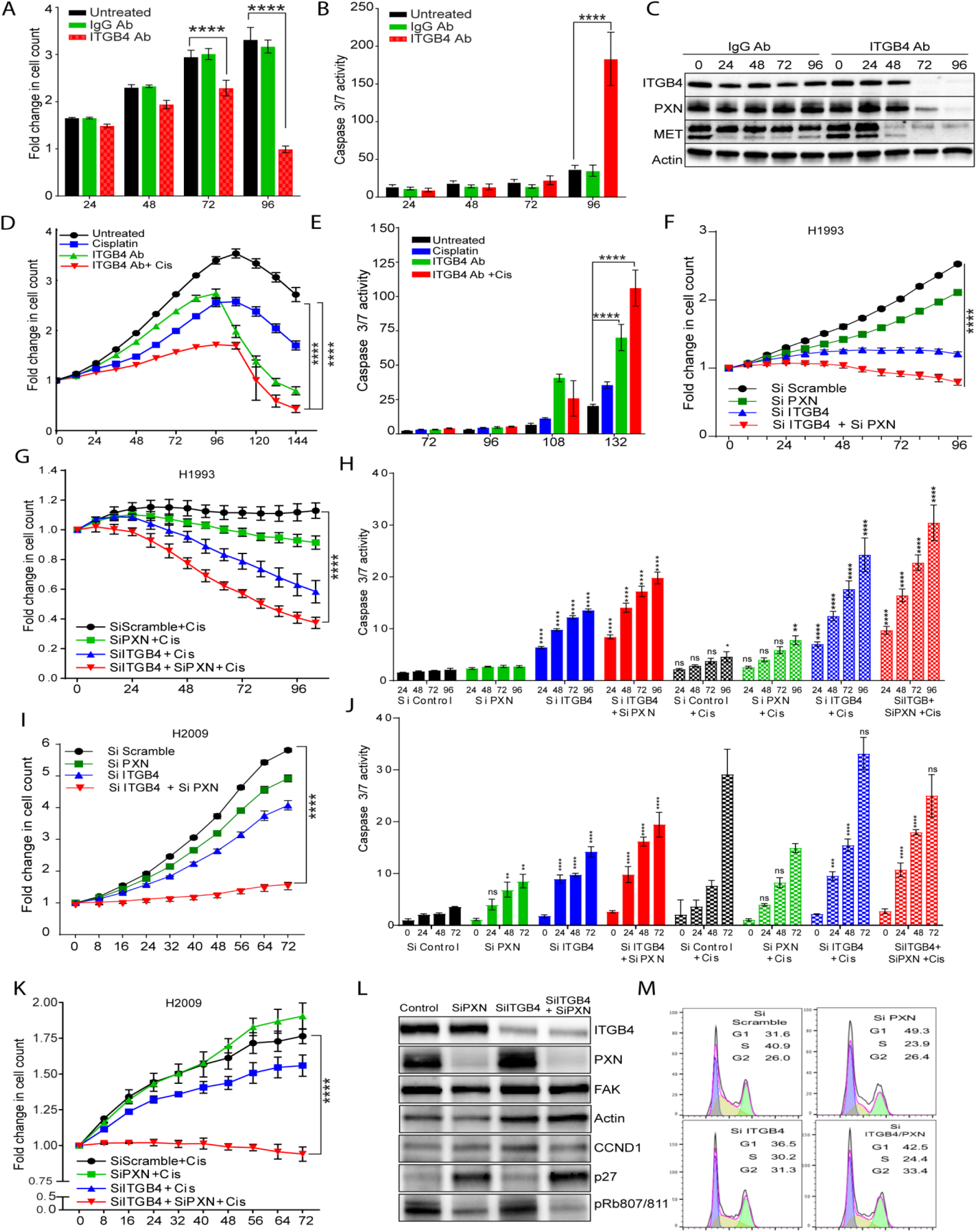
Neutralizing ITGB4 with an antibody or knocking down both PXN and ITGB4 has a synergistic effect on attenuating cisplatin resistance. H1993 cells were treated with an antibody (10 μg/ml) targeting the ITGB4 extracellular epitope in an ultra-low attachment 96-well plate for 6 h then transferred to a tissue culture-treated 96-well plate and allowed to attach overnight. **A)** and **B)** After 96 h, proliferation was inhibited, and apoptosis was significantly induced in cells treated with the ITGB4 antibody (red) compared to untreated (black) and cells treated with an IgG antibody (green) used as a control. (****p<0.0001 Two-way ANOVA, multiple comparison) **C)** Immunoblotting confirmed decreased expression of ITGB4, PXN, and MET with ITGB4 antibody treatment after 48 h compared to IgG antibody control. **D)** and **E)** ITGB4 antibody (10 μg/ml) treatment in combination with a lower dose of cisplatin (2.5 μM instead of 10 μM) exhibited a synergistic effect on inhibition of proliferation and increased caspase activity. (****p<0.0001 Two-way ANOVA) **F)** Double knockdown of both PXN and ITGB4 (red) in H1993 cells had a synergistic effect on inhibiting proliferation compared to single knockdown of either gene (green/blue) and control (black). **G)** Adding 10 μM cisplatin to the double knockdown cells (red) had an even greater effect on inhibiting proliferation compared to single knockdown of either gene (green/blue) and control (black) treated with cisplatin. (****p<0.0001 Two-way ANOVA) **H)** ITGB4 knockdown and double knockdown of PXN and ITGB4 induced apoptosis in H1993 and rendered cells more prone to toxic effects of cisplatin. (****p<0.0001 Two-way ANOVA) **I)** Double knockdown of both PXN and ITGB4 (red) in H2009 cells also had a synergistic effect on inhibition of proliferation compared to single knockdown of either gene (green/blue) and control (black). (****p<0.0001 Two-way ANOVA) **J)** and **K)** Double knockdown of PXN and ITGB4 also induced strong apoptosis and in combination with cisplatin, proliferation was greatly inhibited and caspase activity increased at an earlier time point of 24 h. (****p<0.0001 Two-way ANOVA) **L)** Immunoblotting confirmed siRNA-mediated knockdown of PXN and ITGB4. PXN knockdown alone increased expression of p27 and decreased levels of phospho-Rb (S807/811), indicating cell cycle arrest. **M)** Cell cycle analysis revealed that knocking down PXN induced G1-S arrest whereas knocking down ITGB4 arrested cells in G2-M. Double knockdown of PXN and ITGB4 arrested cells in G1-S and G2-M.

Next, we investigated the effect of combining the antibody and cisplatin treatments to determine potential synergy between the two modalities. Indeed, combining the two treatments showed an additive effect on cell proliferation and caspase-3/7 activity compared to either treatment alone. Furthermore, we also observed a synergistic effect even when a much lower dose (2.5 μM instead of 10 μM) of cisplatin was used (**Fig. 3D and E**), suggesting that tumors expressing high ITGB4 can be treated with anti-ITGB4 antibody in combination with a low dose of cisplatin, which in turn can reduce the undesirable toxicity associated with cisplatin.

Since cisplatin treatment caused changes in expression of FAK and PXN but not ITGB4 (**Fig. 2E**), and all three proteins are required for FA complex formation and activation, we asked if PXN knockdown alone can induce similar phenotype or PXN and ITGB4 double knockdown can synergistically change cell proliferation and survival rates. In H1993 cells, PXN knockdown inhibited cell proliferation by 15% compared to 50% inhibition by ITGB4 knockdown and 70% inhibition caused by double knockdown (**Fig 3F**). Treating H1993 single and double knockdown cells with cisplatin induced cytotoxic effect; a drop in the fold change of cell count by 40% and 60%, respectively was observed compared to control (scramble siRNA) cells treated with cisplatin. However, cisplatin treatments on PXN knockdown cells had a weaker effect (**Fig. 3G**). We further tested the effect of knockdown on apoptosis and found that PXN knockdown did not activate caspase-3/7 but had an additive effect when knocked down together with ITGB4. Further, knocking down ITGB4 alone or together with PXN made the cells more prone to cisplatin toxicity by inducing apoptosis (**Fig. 3H**). In case of the H2009 cells, double knockdown inhibited cell proliferation significantly, indicating a cytostatic effect compared to scramble knockdown (**Fig. 3I**). We observed a similar effect when these double knockdown cells were treated with cisplatin, unlike the cytotoxic effect observed in H1993 cells (**Fig. 3K**). Again, double knockdown or ITGB4 single knockdown induced apoptosis significantly within 24 h of seeding compared to PXN or scramble knockdown. Addition of cisplatin further sensitized these cells as seen by an increase in apoptosis at 24 h and 48 h. (**Fig. 3J**). Immunoblotting analysis confirmed knockdown of both ITGB4 and PXN (**Fig. 3L**). In addition we observed that PXN knockdown induced expression of tumor suppressor p27, with simultaneous reduction in phosphorylation of the retinoblastoma protein Rb1, suggesting cell cycle inhibition. Indeed, cell cycle analysis after knocking down PXN increased G1 population by 20% indicating slowing of G1-S transition, whereas knocking down ITGB4 resulted in partial decrease of S phase population and accumulation in G1 as well as G2 population. In addition, ITGB4 and PXN double knockdown increased further the population of both G1 and G2, indicating cell cycle halt at both G1-S and G2-M phases (**Fig. 3M**), which may explain the synergistic phenotype.

### Knocking down PXN or ITGB4 disrupts spheroid growth

To mimic conditions closer to *in vivo*, we discerned the effect of knocking down PXN or ITGB4 in spheroids (3D) formed from the cell lines used in 2D culture. H2009 cells engineered to express RFP (mKate2) were first transfected with scrambled or PXN/ITGB4-specific siRNA and then seeded in an ultra-low attachment round bottom plate for generating spheroids. Within 4 h of seeding, we observed spheroid formation in wells seeded with cells that were transfected with the control siRNA but not in the cells transfected with ITGB4 and PXN siRNA (**Supplementary Fig. 3G, Supplementary video 9 to 12**). The double knockdown cells took longer time for forming spheroids. Once they formed spheroids, they were monitored for 5 days using IncuCyte live cell imaging system (**Fig 4A**). The spheroid growth and survival were quantitated by monitoring dye intensity and area in real time. We observed a reduction in spheroid area in double knockdown cells by >40% compared to that seen in the spheroids formed by cells transfected with the scrambled siRNA (**Fig. 4B**). Again, spheroids formed using the cells with the double knockdown showed decline in red fluorescence intensity and by day 4, the intensity was only ∼ 30% of the control (**Fig. 4C**). These data suggest that the cell-to-cell interaction, which is essential for the spheroid formation, was disrupted by ITGB4 and PXN knockdown and thus, the observed decrease in spheroid integrity. Additionally, we treated the single and double knockdown spheroids with cisplatin and observed that double knockdown spheroids were most sensitive followed by spheroids with only ITGB4 knocked down. However, PXN knocked down spheroids behaved similarly to that of control (**Fig. 4D and E**). By immunoblotting analysis, we confirmed that spheroids continued to have diminished levels of ITGB4 and PXN by 72 h post-Si RNA transfection (**Fig. 4F**). To confirm that the reduction in spheroid size was due to cell death upon treatment with cisplatin, we assayed them for caspase-3/7 activity using the Essen BioScience assay as described above. We observed a 20-fold higher caspase-3/7 activity with double knockdown compared to Si Scramble by day 5 (**Supplementary Fig. 3F**). We also analyzed caspase activity in control spheroids and double knockdown spheroids treated with cisplatin and found that the double knockdown spheroids exhibited a strong synergistic effect (**Fig. 4G**). Considered together, the data from the 2D and 3D culture experiments revealed that the interaction between PXN and ITGB4 is important for modulating cisplatin resistance in these LUAD cell lines.

**Figure 4.**
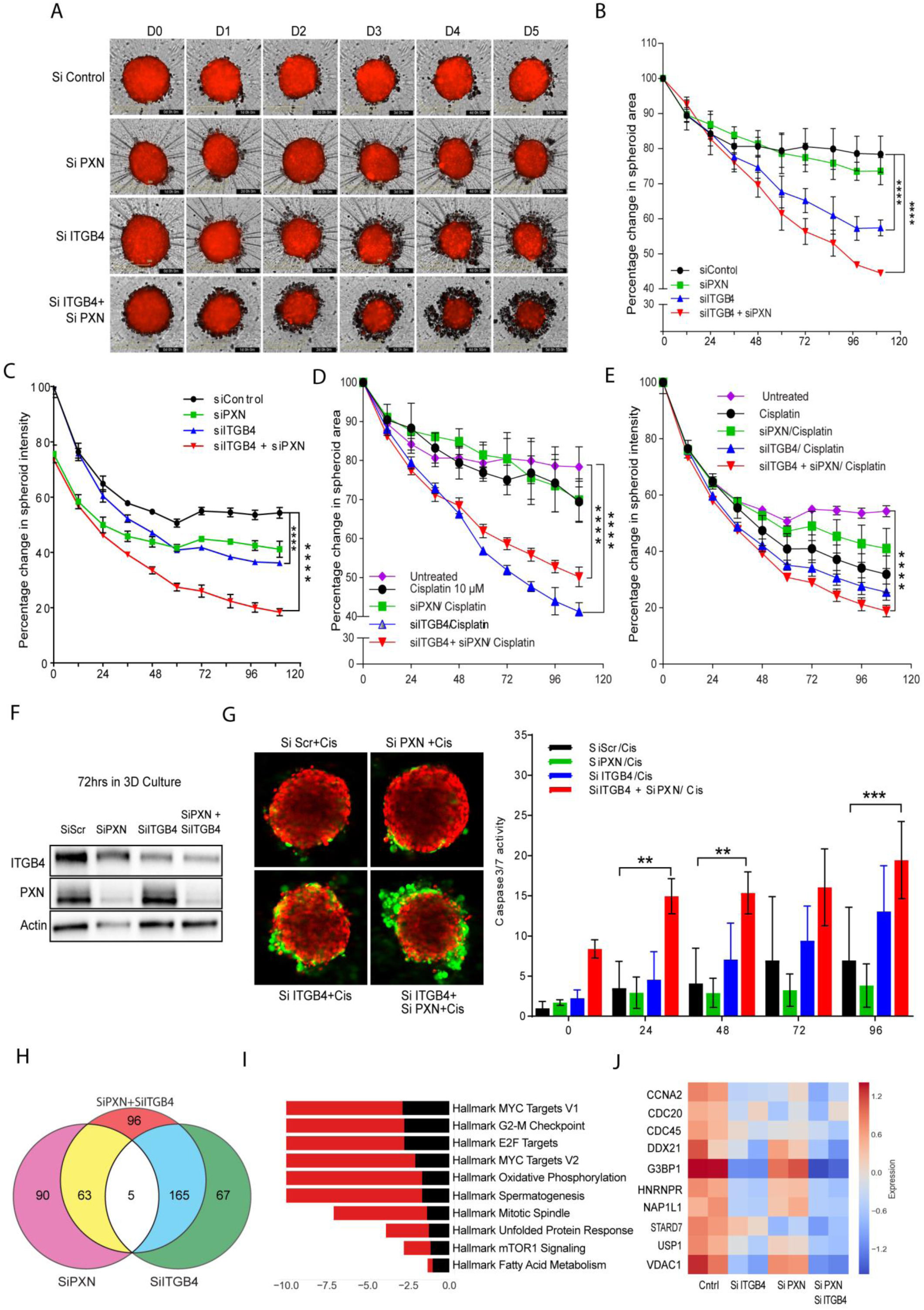
Both PXN and ITGB4 are necessary for spheroid viability and their signaling pathways converge in cisplatin resistance. H2009 cells expressing RFP were transfected with siRNA, seeded in a 96-well ultra-low attachment plate (5000 cells/well), and allowed to form a compact spheroid overnight. **A)** Images acquired by the IncuCyte Live Cell Imaging System showed spheroids with single knockdown to start disintegrating by Day 3 and even earlier for double knockdown spheroids (Day 1). **B)** and **C)** To quantitate spheroid viability, red fluorescence area and intensity were measured. Both parameters showed that double knockdown had a synergistic effect on attenuating spheroid viability. (****p<0.0001 Two-way ANOVA) **D)** and **E)** Spheroids with ITGB4 single knockdown and PXN/ITGB4 double knockdown were sensitized to cisplatin (10 μM) treatment indicated by a decrease in red fluorescence area and intensity. (****p<0.0001 Two-way ANOVA) **F)** Immunobloting confirmed PXN and ITGB4 siRNA-mediated knockdown is still present after 72 h in 3D culture. **G)** Double knockdown spheroids treated with cisplatin had the greatest cytotoxic effect indicated by the green fluorescence in the confocal images acquired by a Zeiss LSM 880 microscope. (**p<0.002, ***p<0.0009 Two-way ANOVA) **H)** Total RNA was extracted from single and double knockdown cells 48 h after siRNA transfection. Total RNASeq revealed the number of genes downregulated with each single knockdown of PXN and ITGB4 and when both genes are knocked down simultaneously. **I)** Bar diagram showing the major Hallmark pathways affected by the double knockdown. The top 10 pathways were arranged in descending order of their enrichment score. **J)** Heat map representation of top 10 genes that belong to hallmark MYC target V1 pathways were analyzed after single or double knockdown. The heat map is representation of two experimental repeats.

### RNAseq analysis suggests that multiple pathways converge in cisplatin resistance

To identify the effect of ITGB4/PXN knockdown on the expression of genes involved in signaling, we determined the changes in global gene expression patterns using RNAseq. RNA was extracted from both single and double knockdown cells 48 h post siRNA transfection and total RNAseq was performed as described in the Methods. In all, 30 million reads were analyzed for each condition. In all, 237 genes were downregulated when ITGB4 was knocked down and 158 genes were downregulated upon knocking down PXN. In the case of the double knockdown, we identified 329 genes that were downregulated (**Fig. 4H**). Interestingly, 5 genes ARL6IP1, GPR160, IFIT1, KIF14, and TSN were common to both single and double knockdown samples (**Supplementary Fig. 4A**). Of these, ARL6IP1 is known to have anti-apoptotic role and KIF14 is known to control cell division, cell cycle progression, and apoptosis.

Comparative GSE analysis of the RNAseq data between control and double knockdown showed significant enrichment of genes regulated by MYC (**Fig. 4I**), genes associated with G2-M checkpoint, and genes regulated by the E2-F transcription factor required for cell cycle progression (**Supplementary Fig. 4B**). Myc is known to induce tumorigenesis by regulating the expression of genes required for suppressing tumor senescence, activating tumor stemness, modifying tumor microenvironment, inducing angiogenesis and suppressing immune activation. In murine models, Myc knockout is enough to contain and eradicate KRAS-driven lung cancer (Soucek et al). Therefore, we determined the effect of ITGB4 and PXN knockdown in regulating MYC targets. The top 10 genes contributing significantly to down regulation of the MYC pathways were interrogated (**Fig. 4J**). Differential expression analysis of RNAseq revealed a list of 96 unique genes which were downregulated only in double knockdown cells. DAVID analysis performed on these 96 genes indicated that they are involved in regulation of transcription, G2/M transition, cell adhesion, and TGF-beta signaling (**Supplementary Table 3**). DAVID analysis of the 206 genes that were downregulated in double knockdown as well as PXN or ITGB4 single knockdown are listed **Supplementary Table 4**.

### ITGB4 and PXN maintain genomic stability and mitochondrial function

The Hallmark pathways represent specific well-defined biological processes and exhibit coherent expression. G3BP1, USP1, and VDAC1 were the top 3 downregulated genes constituting the MYC1 Targets V1 Hallmark pathway. To validate the RNAseq data, we performed western blotting and qPCR experiments. These results confirmed that knocking down either PXN or ITGB4 resulted in the decreased expression of all 3 genes but in double knockdown the decrease in expression becomes more cumulative at both protein (**Fig. 5A**) as well as RNA level (**Supplementary Fig. 4C**). Furthermore, we also analyzed the expression of these genes in lung TCGA data set to determine the expression of USP1 and VDAC1 in tumor and normal samples (**Fig 5B**).

**Figure 5.**
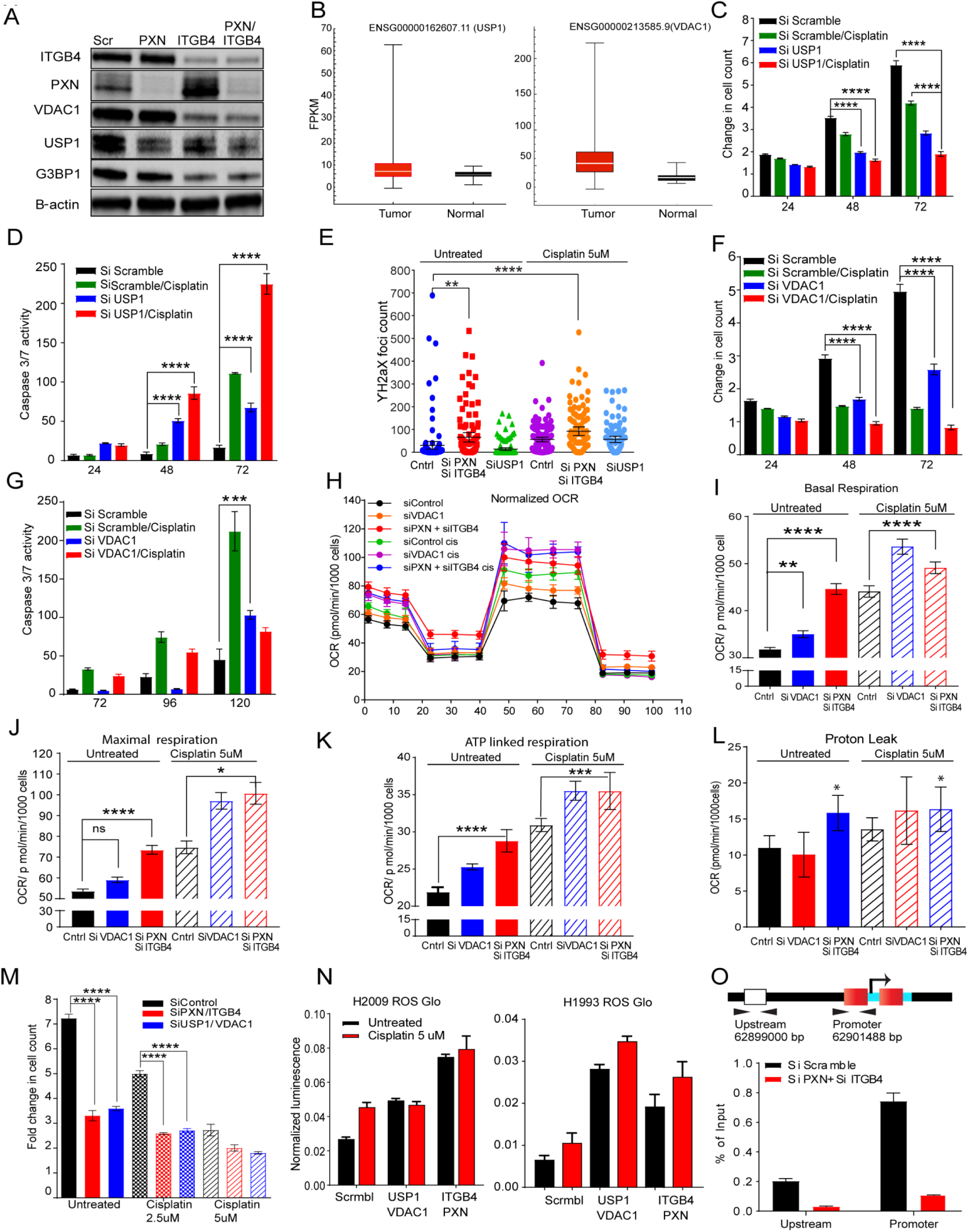
Both PXN and ITGB4 are involved in genomic stability and mitochondrial function. **A)** Immunobloting confirmed decreased expression of top 3 genes (G3BP1, USP1, and VDAC1) downregulated in MYC pathway after knockdown of PXN and/or ITGB4. **B)** Gene expression profiles of LUAD patients were extracted from TCGA database and expression of USP1 and VDAC1 were higher compared to normal tissue. **C)** and **D)** In H2009 cells, knocking down USP1 (blue) attenuated proliferation, induced apoptosis, and sensitized cells to a lower dose of cisplatin (2 μM) (red). (****p<0.0001 Two-way ANOVA) **E)** After knocking down PXN/ITGB4 and USP1, γH2AX foci were detected via immunofluorescence, imaged with confocal microscopy, and counted with QuPath image analysis software. Knockdown exhibited greater number of cells with higher γH2AX foci counts. Added cisplatin increased number of detected γH2AX foci. (*p=0.04, **p=0.004, ***p,<0.0001) **F)** and **G)** Similarly, knocking down VDAC1 (blue) inhibited cell proliferation by 50% within 72hrs, but could induce apoptosis only at later time point. Cisplatin (red) addition has an additive effect to VDAC1 knockdown in inhibiting proliferation. **H)** – **J)** Using the Seahorse XF Analyzer, cellular metabolic activity was measured with knockdown and in presence of cisplatin. Knocking down PXN/ITGB4 or VDAC1 increased basal **(I)** and maximal **(J)** mitochondrial oxygen consumption rate per 1000 cells. (**p=0.0029, ****p<0.0001 One-way ANOVA) **K)** and **L)** ATP-linked respiration **(K)** and proton leak **(L)** also showed an increase in the same knockdown cells. (* p=0.01 One-way ANOVA) **M)** Double knockdown of PXN/ITGB4 (red) and USP1/VDAC1 (blue) inhibited proliferation to similar extent whether in the absence or presence of cisplatin. (****p<0.0001 Two-way ANOVA) **N)** Reactive oxygen species (ROS) production was measured using the ROS-Glo™ H2O2 Assay. Double knockdown of PXN/ITGB4 and USP1/VDAC1 induced higher levels of ROS compared to control. **O)** ChIP was performed with an acetylated H3K27 antibody 72 h after siRNA-mediated knockdown of PXN/ITGB4. With the knockdown, H3K27 acetylation at the promoter region of USP1 was greatly reduced compared to that of an upstream region, indicating ITGB4 and PXN role in USP1 transcriptional activation.

We next asked whether knocking down USP1, G3BP1 or VDAC1 gene would recapitulate the phenotype of the ITGB4/PXN double knockdown. To this end, we used 3 siRNA constructs for each gene USP1, VDAC1, and G3BP1 (OriGene). 10 nM of each siRNA was transfected in the H2009 cells and the optimal siRNA construct was selected by correlating knockdown efficiency and phenotype (fold change in cell proliferation) (**Supplementary Fig. 4D**). G3BP1 siRNA knockdown inhibited its expression but did not inhibit cell proliferation or induce caspase activity (**Supplementary Fig. 4D and E**). For USP1, siRNA sequence B and for VDAC1 siRNA sequence A were selected for all downstream experiments.

USP1 knockdown inhibited cell proliferation by more than 2-fold within 72 h of seeding with simultaneous increase in caspase activity. Furthermore, addition of a lower dose of cisplatin (2.5 μM) to USP1 knocked down H2009 cells decreased their proliferation by 3-fold (**Fig. 5C**) and increased apoptosis by 200-fold within 72 h (**Fig. 5D**). We also tested the sensitization of H2009 and H1993 cells to the USP1-selective inhibitor ML323 (Qin et al). We treated cells with ML323, 5 μM and 10 μM for 3 days but did not see any significant change in the cell proliferation rates (**Supplementary Fig. 4F**). These data indicate that H2009 and H1993 cell survival and proliferation is dependent on USP1 protein level. ML323, a USP1 inhibitor known to disrupt the USP1-UAF interaction and may not decrease USP1 protein level and hence, could not reproduce the phenotype of USP1 knockdown. Furthermore, USP1 is a deubiquitinase known to increase DNA repair activity in cisplatin-treated cells (Iraia et al). Therefore, we sought to measure changes in the γH2AX foci as a measure of DNA damage in the double knockdown or USP1 knockdown cells. We observed that double knockdown cells have significantly higher γH2AX foci counts compared to control or USP1 knockdown. Addition of cisplatin in conjunction with USP1 knockdown further increased the extent of DNA damage (**Fig. 5E**).

On the other hand, VDAC1 is an ion channel pump located in the mitochondrial and plasma membranes. Cox regression analysis on VDAC1 expression identified it as an independent factor with significant prognostic value (p<0.0001) for worse overall survival (Calire eta al). Therefore, we determined the significance of VDAC1 knockdown in our experimental set up. Knocking down VDAC1 led to a decrease in cell proliferation by 50% within 72 h. However, this did not contribute to apoptosis induction at an early time point even in the presence of cisplatin. Although at a later time point (120 h), we observed an increase in apoptosis by 2-fold (**Fig. 5F and G**). To discern the role of VDAC1 in mitochondrial function, we knocked down VDAC1 as well as PXN and ITGB4 in H2009 cells and analyzed mitochondrial respiration in absence and presence of cisplatin using the Seahorse XF Analyzer (Agilent, Santa Clara, CA). The raw data was normalized to the live cell count. Knocking down VDAC1 or ITGB4/PXN changed the oxygen consumption rate (OCR) of cells, which is measured by the basal as well as maximal mitochondrial OCR compared to the control, and these rates were further increased in presence of cisplatin (**Fig. 5 H, I, and J**).

Oxidative phosphorylation is an efficient but slow process of ATP production in comparison to the glycolytic pathway and is more preferably used by tumor cells. Pyruvate generated during glycolysis is oxidized by mitochondria for oxidative phosphorylation leading to mitochondrial ATP production, which is required for ATP-linked respiration (Ajit et al). Treating cells with Oligomycin, which inhibits the electron transport chain and blocks mitochondrial ATP generation, can inhibit ATP-linked respiration. On one hand, an increase in ATP-linked respiration is indicative of the availability of more substrates like pyruvate to drive oxidative phosphorylation. On the other hand, it also suggests an increase in ATP demand of the cell. Therefore, we calculated ATP-linked respiration and observed an increase in the linked respiration in double knockdown cells which increased further upon addition of cisplatin. Control (scramble siRNA treated) cells in the presence of cisplatin showed increased ATP-linked respiration which is indicative of a stress-induced increase in ATP demand. The same increase in ATP demand was also seen in these cells upon VDAC1 knockdown or double knockdown (**Fig. 5K**).

Mitochondria maintain a proton motive potential for ATP generation under ideal conditions. A proton leak can affect the membrane potential and consequently, a decrease in ATP production. In order to maintain the proton motive force intact, mitochondria increase their respiration and oxygen consumption rate as measured by increase in basal and maximal respiration. We observed an increase in the proton leak for ITGB4 and PXN double knockdown which may explain their increase in respiration to compensate for the loss in membrane potential (**Fig. 5L**). Further, respiratory capacity is measured by considering the ratio of maximal respiration compared to basal respiration, which does not show much difference across samples (**Supplementary Fig 4G**).

We also checked the possibility of decreased expression of ITGB4 and PXN upon knocking down USP1 or VDAC1 but did not observe any change (**Supplementary Fig. 4H**).Taken together, these data suggest that USP1 and VDAC1 are downstream of PXN and ITGB4 and thus, knocking down USP1 and VDAC1 should recapitulate the phenotype of PXN and ITGB4. Therefore, we transfected H2009 cell line with 10 nM of siRNA against ITGB4 and PXN or siRNA against USP1 and VDAC1 and compared changes in proliferation rates. Cell proliferation was reduced by 50% after knocking down either of the combination. Addition of cisplatin to either combination had an additive effect in inhibiting cell proliferation. Interestingly, there was no significant difference between either gene combinations (**Fig. 5M**).

Irradiation is known to increase mitochondrial respiration causing an increase in generation of mitochondrial ROS which eventually induces apoptosis (Tohru Yamamori et al). Therefore, we asked if increased mitochondrial respiration leads to increased reactive oxygen species (ROS) generation in the double knockdown cells. We estimated the production of ROS using ROS-Glo Kit from Promega in two cisplatin-resistant cell lines by knocking down USP1/VDAC1 or ITGB4/PXN. We observed an increase in the generation of ROS by knocking down either ITGB4/PXN or USP1/VDAC1 compared to control (scramble siRNA) knockdown (**Fig. 5N**). Addition of cisplatin to control cells increased ROS production but had no significant effect on the double knockdown cells. In previous experiments we did not observed any change in the expression of PXN and ITGB4 on knocking down either USP1 or VDAC1. Again we did not observe any change in PXN and ITGB4 expression in USP1 and VDAC1 double knockdown cells indicating the regulation of USP1 and VDAC1 by PXN and ITGB4 is unidirectional. (**Supplementary Fig. 4I**)

Taken together, the data suggests that ITGB4 and PXN regulate USP1 and VDAC1 expression, which plays a critical role in the tumor proliferation and sensitivity to cisplatin. Knocking down either ITGB4/PXN or USP1/VDAC1 increases mitochondrial oxygen consumption rate, leading to increase in ROS generation which eventually leads to DNA damage. The induction of DNA damage was cumulative when cisplatin was added to these cells. To repair this extensive DNA damage, cells need to activate the DNA repair mechanism like USP1 activation, which were abrogated by ITGB4 and PXN double knockdown.

Finally, since double knockdown silenced USP1 expression alluding to possible transcriptional regulation, we analyzed the upstream sequence of USP1 promoter region using Genome Browser and identified two potential acetylated histone H3K27 binding sites. To determine the functional significance of these sites, we knocked down both PXN and ITGB4 and after 72 h, performed chromatin immunoprecipitation (ChIP) using an anti-acetylated histone H3K27 antibody from Diagenode. In the double knockdown samples, we observed reduced binding of acetylated H3K27 at the promoter, suggesting that USP1 expression is induced by hyper acetylation (**Fig. 5O**). Of note, the data also suggest that ITGB4 and PXN involvement is not limited to migration and invasion; they may also be involved in controlling the epigenetic landscape of lung cancer. However, additional studies are needed at the global level along with other histone acetylation forms.

### Biochemical data demonstrate direct interaction between PXN and ITGB4

From the foregoing, it follows that PXN and ITGB4 interact with each other and also with FAK and that this interaction is important in cisplatin resistance. To determine these interactions, we performed co-immunoprecipitation experiments with an ITGB4 antibody as well as with FAK or PXN antibodies in separate experiments. Immunoblotting with ITGA6 antibody served as a positive control as it is known to interact with ITGB4 (Aydin et al). To our surprise however, we were unable to detect the ITGA6 protein in H2009 and H1993 cell lines. However, the pulldown showed the presence of ITGA7 (**Fig 6A**). Furthermore, the immunoprecipitation data indicated a stronger interaction between FAK and ITGB4 compared to PXN and ITGB4, suggesting that FAK interaction with ITGB4 is PXN-dependent or vice versa. To test this possibility, we knocked down PXN and repeated immunoprecipitation with the ITGB4 antibody. This experiment demonstrated that indeed, FAK interacts with ITGB4 irrespective of PXN (**Fig. 6B**). We next performed immunoprecipitations using lysates from cells treated with cisplatin for 48 h to determine if cisplatin induced any change in the interaction of ITGB4, FAK, and PXN (**Fig. 6C and D**). By immunoprecipitating ITGB4 as well as PXN, we observed no change in the interactions among FAK, PXN, and ITGB4 in presence of cisplatin, which again indicated the involvement the FA complex in imparting cisplatin resistance. Considered together, the co-immunoprecipitation data suggested that disrupting interaction between PXN, ITGB4, and FAK in cisplatin-resistant cells could make them more susceptible to cisplatin.

**Figure 6.**
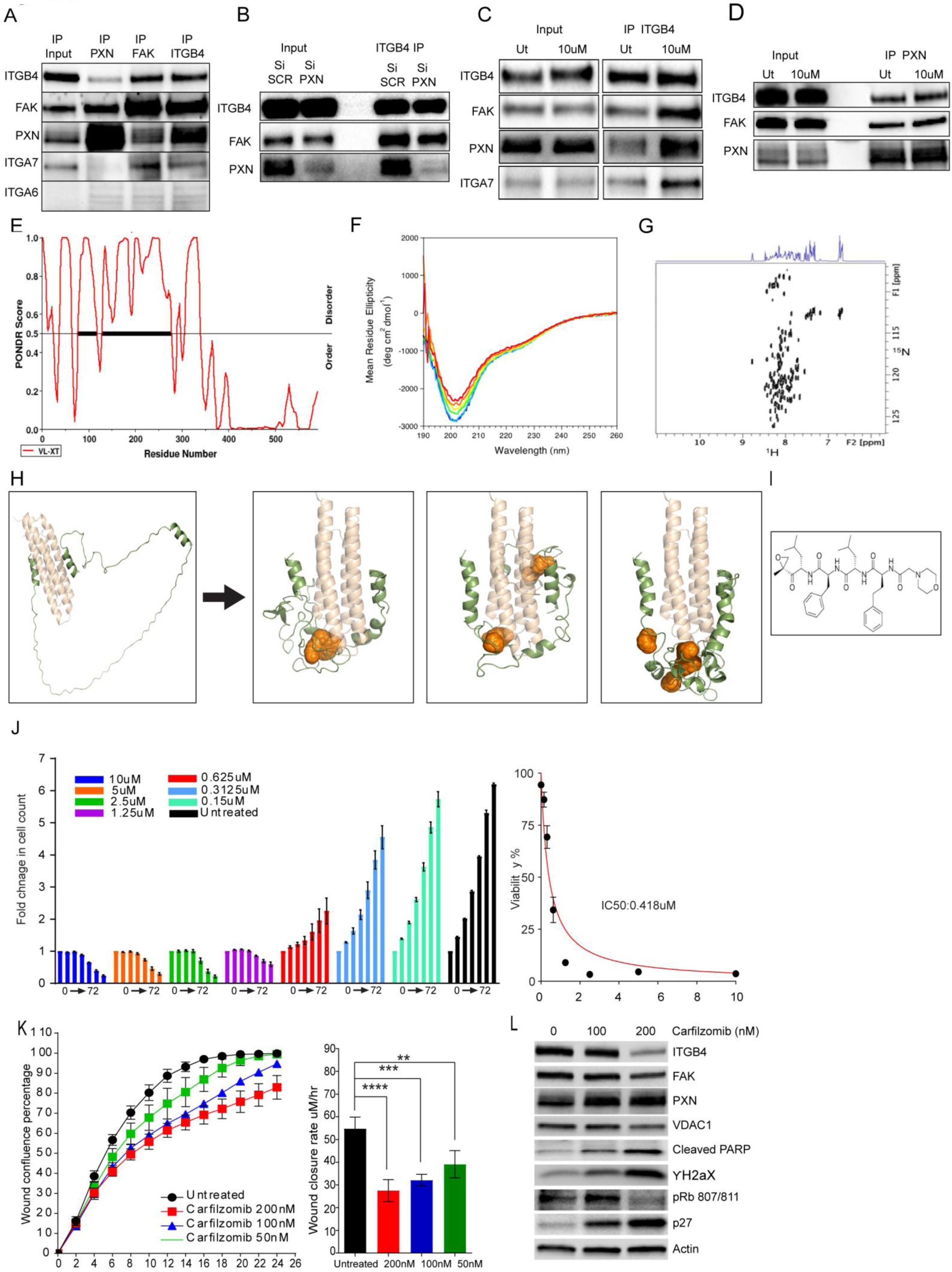
PXN is intrinsically disordered and its interaction with ITGB4 can be disrupted by an FDA-approved compound. **A)** Co-immunoprecipitation (co-IP) of H2009 whole cell lysate with an ITGB4 antibody showing FAK, PXN, and ITGA7 interact with ITGB4. Reverse co-IP with FAK and PXN antibody showing ITGB4 binds to FAK and PXN. **B)** Knocking down PXN did not affect the interaction between FAK and PXN. **B)** and **D)** Treating H2009 cells with 10 μM cisplatin for 48 h did not induce any changes in the interaction between ITGB4, FAK, and PXN. **E)** PONDR prediction algorithm determined that the N-terminal region of PXN to be intrinsically disordered whereas the C-terminal half is highly ordered. **F)** Circular dichroism (CD) spectra were recorded as a function of temperature 10°C to 60°C. Low ellipticity values near the 215-230 nm region indicated that the N-terminal region lacks any secondary structure. **G)** NMR spectroscopy using a two-dimensional ^1^H-^15^N HSQC spectrum was used to analyze the LD2-LD4 region of PXN to confirm that it is indeed disordered. **H)** Using the binding pocket in the N-terminal region of PXN, 1440 FDA-approved drugs were virtually screened. **I)** Molecular structure of carfilzomib. **J)** H2009 cells were treated with an increasing dose of carfilzomib to determine IC50. **K)** Scratch wound assay using H2009 cells treated with a sublethal dose of carfilzomib (50-200 nM) showed dose-dependent inhibition of cell migration. (**p-0.002, ***p-0.0001, p-****<0.0001) **L)** Immunoblot of H2009 cells treated with carfilzomib showed reduced expression of ITGB4 and phospho-Rb (S807/811) and increased expression of p27, γH2AX, and cleaved PARP, indicating cell cycle inhibition and cell death.

### The N-terminal region of PXN is intrinsically disordered

Since PXN, especially the LD1-LD5 domain, is implicated in interacting with multiple proteins in the FA complex, we suspected that this region may be intrinsically disordered. Bioinformatics analysis using the PONDR prediction algorithm showed that indeed, the N-terminal half of the molecule, especially the regions connecting the LD domains, is predicted to be significantly disordered while the C-terminal half comprising the LIM domains is predicted as highly ordered (**Fig. 6E**). To experimentally verify the prediction, we employed circular dichroism (CD) and NMR spectroscopy. CD spectra were recorded as a function of temperature from 10° to 60°C. These spectra showed that the N-terminal region of PXN polypeptide chain lacks significant α-helical or β-strand secondary structural elements over this temperature range as evidenced by low ellipticity values in the 215–230 nm region. However, a weak ellipticity minimum at ∼222 nm was observed, suggesting some helical character which could likely correspond to the short helical regions (5-7 amino acids) between the large disordered regions (**Fig. 6F**). To corroborate the CD data, we used NMR. A two-dimensional ^1^H-^15^N HSQC spectrum of the LD2-LD4 region (**Fig. 6G**) confirmed that this domain is indeed disordered. Taken together, the biochemical, computational, and biophysical experiments not only provide good evidence that PXN is an intrinsically disordered protein (IDP) and therefore can interact with multiple partners, but also suggest that the interactions between PXN, ITGB4, and FAK contribute to cisplatin resistance in lung cancer. Consistently, our data demonstrating a strong synergistic effect of the double knockdown underscored the importance of the PXN/ITGB4 complex in drug resistance in these cells.

### FDA approved drugs alleviate cisplatin resistance in LUAD

A virtual screen of a library of 1440 FDA-approved drugs using binding pockets in the N-terminal domain of PXN (**Fig. 6H**) identified several putative hits. The top 40 hits were then rescreened employing a cell-based assay to determine their efficacy using H358 cells. We used the H358 cells because they have high expression of ITGB4/PXN and exhibit intermediate sensitivity to cisplatin. Cells were treated with various concentrations of the drugs (0.15 to 10 μM) for 72 h and the IC50 values were determined using the GraphPad software (**Supplementary Table 5**). We identified 11 potential compounds that showed a significant decrease in cell proliferation with IC50s ranging from 0.8 nM to 1.54 μM. We then used a sublethal dose to determine the effect of these compounds on ITGB4 and PXN expression and caspase activity (**Supplementary Fig. 5A**). Three of the 11 compounds carfilzomib, ixazomib, and CUDC-101 induced apoptosis and caused DNA damage measured by γH2AX immunblotting and decrease in ITGB4 expression. We repeated the drug assay using the 11 compounds on H2009 cell line using a range from 10 μM to 0.08 μM. The 0.08 μM of carfilzomib is highly potent to inhibit cell proliferation as compared to the other 10 compounds (**Supplementary Fig. 5B**). CCK-8 viability assay was done on the cells treated with 0.08 μM of drugs, and again carfilzomib emerged as a strong inhibitor of cell viability (**Supplementary Fig 5C**). Further immunoblotting done on the H2009 cells at a sublethal dose using all the 11 compounds showed degradation or loss of ITGB4 and PXN in the cells treated with carfilzomib only (**Supplementary Fig. 5D**). We further tested the effect of of top3 drugs carfilzomib, ixazomib, and CUDC-101on H2009 and H1993 cell lines. Of these, carfilzomib was the most efficacious with an IC50 of 0.418 μM (**Fig. 6I and J**) and 1.7 μM (**Supplementary Fig. 5E**) in H2009 and H1993, respectively. Furthermore, we also confirmed significant inhibition of wound healing and reduction in wound closure rate in H2009 cells when treated with sublethal doses of 200, 100, or 50 nM of carfilzomib (**Fig. 6K**). Ixazomib and CUDC-101 on the other hand, were less effective than carfilzomib in inhibiting wound healing (**Supplementary Fig. 5G**). Finally, immunoblotting experiments using cell extracts from H2009 and H1993 cells treated with a sublethal dose of carfilzomib showed a reduction in ITGB4 expression together with an increase in the levels of p27, γH2AX, and cleaved PARP (**Fig. 6L and Supplementary Fig. 5F**).

### Carfilzomib inhibited patient-derived organoids better than cisplatin

The experiments described above were done using 2D monolayer cultures. To discern the effect of the drug in 3D cultures, we employed H2009 spheroids. The spheroids were treated with different doses (150-600 nM) for 72 h and spheroid integrity was analyzed by measuring the red and green intensities using confocal microscopy. Consistent with the 2D data, a significant increase in caspase activity and decrease in spheroid growth was observed. The confocal images taken 72 h post-treatment showed intact spheroids in the untreated condition but disintegrated spheroids with high green fluorescence (caspase activity) in the presence of carfilzomib (**Fig. 7A-C**).

**Figure 7.**
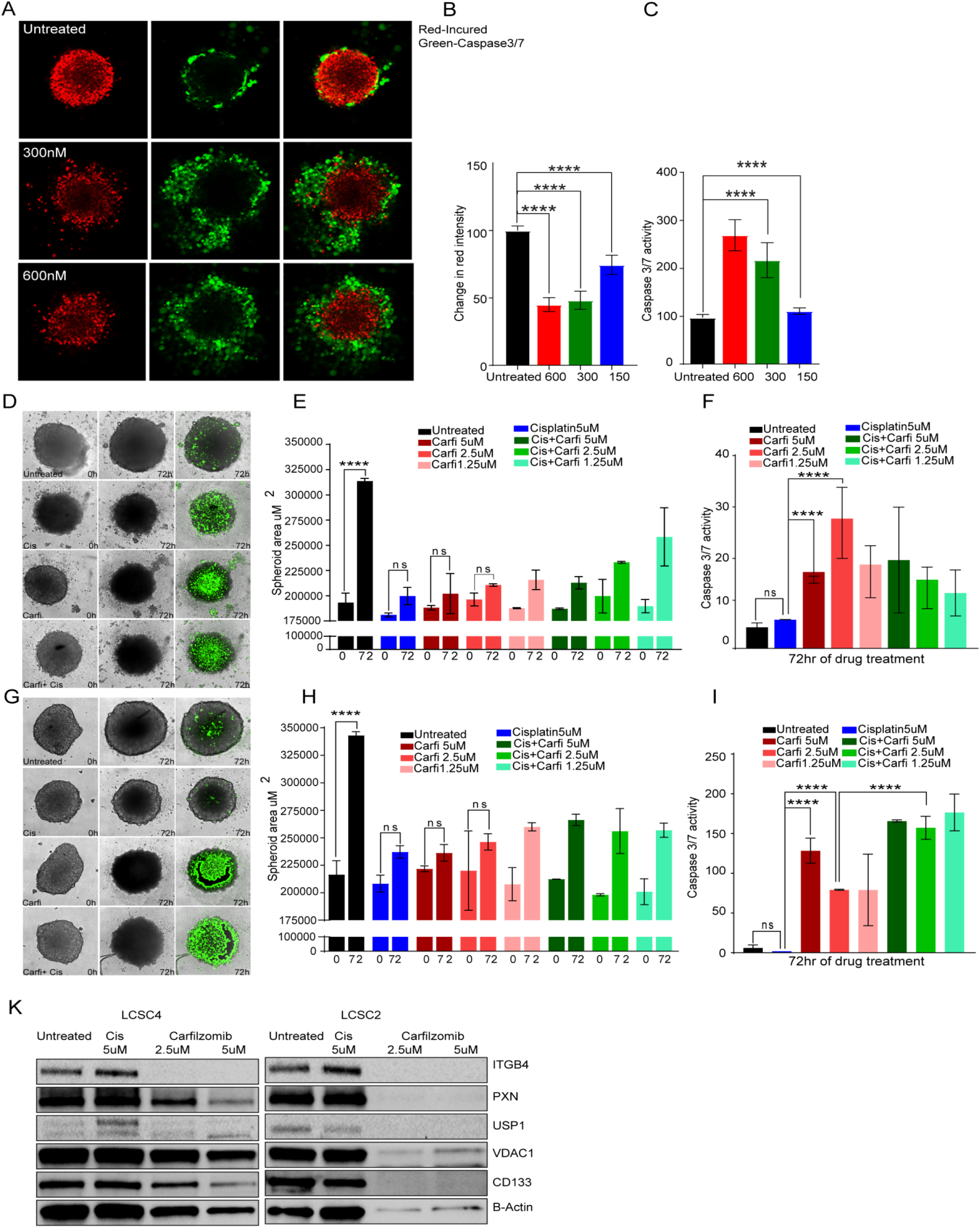
Carfilzomib is more effective than cisplatin in 3D spheroids and patient-derived organoids. **A)** Confocal images taken of H2009 spheroids treated with 300 nM and 600 nM carfilzomib for 72 h depicted disintegration and induction of caspase activity marked by green fluorescence. **B)** and **C)** Carfilzomib treatment of H2009 spheroids reduced viability of spheroids as quantitated by lower red fluorescence intensity **(B)** and increased caspase activity **(C)** in a dose-dependent manner. (****p<0.0001 Ordinary one-way ANOVA) **D – I)** Surgical samples from lung cancer patients were obtained and cultured to form organoids (5000 cells/well). In one patient sample (LCSC4) **(D)**, cisplatin (5 μM) alone and carfilzomib (2.5 μM) alone decreased the size of the organoid **(E)**, but there was a 5-fold greater induction of apoptosis with carfilzomib compared to cisplatin in reference to cisplatin, (****p<0.0001 Two-way ANOVA) **(F)**. The combination of carfilzomib and cisplatin did not have an additive effect. In another patient sample (LCSC2) **(G)**, cisplatin and carfilzomib decreased the spheroid area **(H).** Cisplatin alone did not induce any caspase activity, but carfilzomib induced a ∼50-fold increase in apoptosis. (****p<0.0001 Two-way ANOVA) **(I)**. Caspase activity was enhanced with the combination of both drugs. (****p<0.0001 Two-way ANOVA) **J)** These patient-derived organoids were treated with cisplatin and carfilzomib at indicated concentrations and collected after 48 h for immunoblot. Cisplatin treatment had a minimal effect whereas carfilzomib had a more toxic effect on cells marked by decreased expression of ITGB4, PXN, USP1, VDAC1, and CD133.

Next, we determined drug efficacy in patient tumor tissue-derived organoids form two NSCLC adenocarcinoma patients. The organoids were named as lung cancer stem cell (LCSC) because of the expression of the stem cell markers (Xinwei et al). Approximately 5000 cells/sample from 2 surgical samples were seeded in an ultra-low attachment plate and incubated at 37°C. Once organoids formed, carfilzomib and caspase assay dye were added, and the organoids were observed in real time using the IncuCyte Live Cell Analysis System. As expected, in one of the patient-derived organoids, cisplatin treatment alone decreased spheroid area and increased apoptosis as measured by green intensity (**Fig. 7D**). Furthermore, treating with carfilzomib also decreased spheroid growth and increased apoptosis. Indeed, by 72 h carfilzomib at a concentration of 2.5 μM induced 5-fold increase in caspase activity compared to cisplatin alone but there was not much change in the spheroid area (**Fig. 7E and F**). - Thus, no additive effect was apparent in this patient sample. However, in the second case, cisplatin did not induce any caspase activity but carfilzomib by itself induced ∼50-fold increase in caspase activity which was further enhanced when the two drugs were combined (**Fig. 7G-I**). We also tested the efficacy of ITGB4 antibody in disrupting organoid growth and inducing apoptosis. After 6 days of treatment, a significant decrease in organoid area was observed, suggesting that ITGB4 antibody treatment significantly affects patient-derived organoid growth compared to untreated or 2.5 µM cisplatin-treated, but could not disrupt the organoid or induce cytotoxicity (**Supplementary Fig. 5H**). Considered together, these data indicated that in patients who are resistant to cisplatin, sensitivity can be enhanced by using carfilzomib.

Next, we treated the organoids with cisplatin alone or carfilzomib at 2.5 µM and 5 µM for 48 h and processed the samples for immunobloting. We observed a decrease in the expression of ITGB4, USP1, PXN and VDAC1 which were found to be associated with cisplatin resistance in the present study (**Fig. 7G**). In addition, we also observed decrease in the expression of CD133 expression, a marker for cancer initiating cells, which are thought to be responsible for acquired drug resistance (Bertolini G et al). Thus, carfilzomib treatment can sensitize the tumor cells as well as the cancer initiating cells, suggesting a potential drug for cisplatin refractory lung cancer cases.

### Mathematical modeling indicates bistability

Analysis of RNAseq data revealed that several microRNAs were also differentially regulated in response to single or double knockdown (**Supplementary Table 6**). In particular, mir-1-3p was upregulated both in ITGB4 single knockdown as well as in the ITGB4/PXN double knockdown by 3.1-fold and 2.54-fold, respectively. Furthermore, it was reported that overexpressing ITGB4 downregulates miR-1-3p (Gerson et al, 2012). Together, these observations suggested that there exists a double negative feedback loop between ITGB4 and miR-1-3p. Therefore, integrating these observations, we developed a mathematical model to simulate the dynamics of ITGB4/miR-1-3p feedback loop (**Fig. 8A**). This model showed the existence of two stable states (phenotypes): one (sensitive) state represented by (low ITGB4, high miR-1-3p) and the other (resistant) state represented by (high ITGB4, low miR-1-3p) (**Fig. 8B**). Moreover, the model suggested it is possible for cells to switch their state under the influence of biological noise or stochasticity; thus, a sensitive cell can behave as resistant and *vice-versa*.

**Figure 8.**
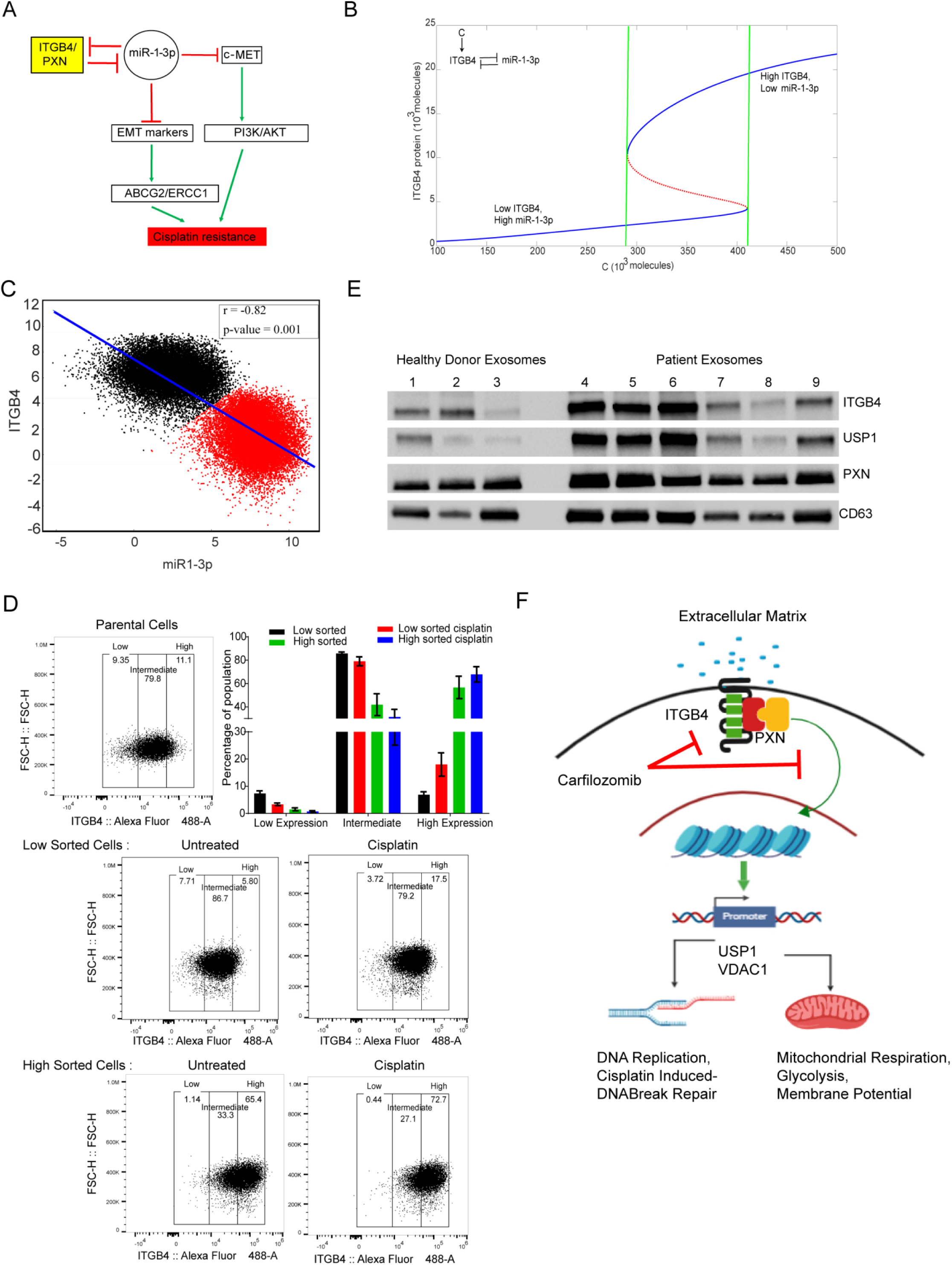
Mathematical modeling suggests that cisplatin resistance can be stochastic and reversible state, and ITGB4 and PXN detected in exosomes represent attractive biomarkers. **A)** A double negative feedback loop between ITGB4 and miR-1-3p leads to bistability. **B)** A mathematical model stimulating the dynamics of ITGB4 and miR-1-3p showed cisplatin resistance to be a reversible state. High ITGB4 and low miR-1-3p render cells to be resistant whereas low ITGB4 and high miR-1-3p represents a more state. **C)** To test this mathematical model, RACIPE algorithm generated an ensemble (n = 100,000) with varying parameter sets then plotted to represent robust dynamical patterns. Results showed that ITGB4 and miR-1-3p exhibit bimodality: two distinct subpopulations of cells that are negatively correlated, reinforcing the previously described negative feedback loop. **D)** H2009 cells were stained with ITGB4 antibody conjugated to Alexa Fluor 488 and sorted based on gates set to high and low ∼10% of ITGB4-expressing population using the FACSAria Fusion instrument. Sorted cells were subsequently cultured for 48 h and then treated with 1 μM cisplatin for 48 h. Then using the Attune NxT Flow Cytometer, equal numbers of cells were stained again and analyzed to determine shifts in population between untreated and treated cells. Low sorted cells treated with cisplatin had a greater cell population that shifted toward higher ITGB4 expression. High sorted cells treated with cisplatin did not undergo significant changes in population compared to untreated. **E)** Human sera (500 μl) were obtained from healthy donors and LUAD patients. Exosomes were isolated, lysed, and analyzed for expression of ITGB4 and PXN by immunoblotting. Compared to that of healthy donors, exosomes of patients had increased expression of ITGB4 and PXN and also USP1, which we identified to be regulated by ITGB4. **F)** Schematic depicting the interaction between ITGB4 and PXN regulating downstream proteins USP1 and VDAC1 at the transcriptional level to coordinate cisplatin resistance.

Next, to investigate whether this co-existence of two states is a robust feature that can be expected from the given network topology, we implemented the algorithm RACIPE (Random Circuit Perturbation) which generates an ensemble of mathematical models with varying parameter sets (Huang et al, 2017). The results from this ensemble (n=100,000) were then plotted together to identify robust dynamical features of the underlying regulatory network; each mathematical model can represent one cell, and this ensemble represents a cell population with varying levels of genetic/phenotypic heterogeneity. RACIPE results for the ITGB4/miR-1-3p feedback loop showed that both ITGB4 and miR-1-3p exhibit bimodality, i.e. two subpopulations (**Fig. 8B and C**). These two subpopulations were also distinctly observed in a scatter plot of ITGB4/miR-1-3p showing negative correlation (r = - 0.82, p <0.001), reinforcing our previous results indicating the existence of two states – high ITGB4, low miR-1-3p and low ITGB4, high miR-1-3p. All in all, these results indicate that ITGB4/miR-1-3p feedback loop enables phenotypic plasticity.(Supplementary Fig. 6A)

To test the predictions made by the model, we treated H2009 cells with cisplatin for 4 days. On day 5, we subjected them to FACS analysis using an ITGB4 antibody. We observed a 15% decrease in ITGB4 low-expressing cells compared to untreated cells, suggesting that the H2009 cells are inherently a mixed population with variable expression of ITGB4. In presence of stress, the cells either increase the expression of ITGB4 or the cells having high ITGB4 get selected (**Supplementary Fig. 6B**). Additionally, we gated and purified 10% of median low ITGB4 expressing cells (Low Sorted) and 10% of median high ITGB4 (High Sorted) subpopulations from H2009 cells, by FACS sorting and cultured them separately. We seeded the sorted cells in 6 independent wells and allowed them to proliferate. After 2 days, 3 out of 6 wells were treated with a sub-lethal dose of cisplatin (1μM) for 2 days. The cells were then counted, and an equal number of cells were stained with ITGB4 antibody and PI and analyzed by Attune NxT Flow Cytometer. Interestingly, we observed that the ITGB4-low cells recreated ITGB4-intermediate and ITGB4-high population while ITGB4-high sorted cells kept on proliferating and increasing the population of cells having highITGB4. The high sorted cells crated an intermediate population of 30% where as the low sorted created a population of 80% similar to the parental cells. Thus indicating cells having intermediate or high ITGB4 expression were more preferentially favored. Finally, we treated these cells with cisplatin and observed a shift in population for the low and intermediate gated cells towards the higher gate, suggesting that (cisplatin) stress favored the population with high expression of ITGB4 (**Fig. 8D**).

### Exosomal ITGB4 and PXN as biomarkers for predicting cisplatin response

Since ITGB4 and PXN are upregulated in cisplatin-resistant cells, we asked if a minimally invasive method could be developed to determine the expression of ITGB4 and PXN in LUAD patients. To this end, we used 500 μl of serum from healthy donors and LUAD patients and isolated exosomes by ultracentrifugation as described in the Methods. Exosomes were lysed and analyzed for ITGB4 and PXN expression by western blotting. CD63 served as an exosomal marker protein. Indeed, we observed that of six patients that were tested, three patients had higher expression of ITGB4 and PXN. Interestingly, USP1, which we identified as being regulated by ITGB4, showed a similar pattern corresponding to ITGB4 (**Fig. 8E**). These data suggest that ITGB4 and PXN represent attractive predictive biomarkers for cisplatin resistance.

## Discussion

It is generally held that the antineoplastic effects of cisplatin are due to its ability to generate unrepairable DNA lesions hence inducing either a permanent proliferative arrest (cellular senescence) or cell death due to apoptosis. The drug enters cells via multiple pathways and forms multiple DNA-platinum adducts which results in dramatic epigenetic and/or genetic alternations. Such changes have been reported to occur in almost every mechanism supporting cell survival, including cell growth-promoting pathways, apoptosis, developmental pathways, DNA damage repair, and endocytosis (Shen et al, 2012; Rocha et al, 2018). Therefore, at a single cell level, the genetic underpinning involved in cisplatin resistance is obvious; however, the exact molecular mechanism(s) underlying the emergence of cisplatin resistance remains poorly understood and a bewildering plethora of targets have been implicated in different cancer types.

On the other hand, drug resistance is also thought to be strongly influenced by intratumoral heterogeneity and changes in the microenvironment (Álvarez-Arenas et al, 2019). Again, while prevailing wisdom advocates that the heterogeneity arises from genetic mutations, and analogous to Darwinian evolution, selection of tumor cells results from the adaptation to the microenvironment (Gerlinger M, Swanton, 2010), it is now increasingly evident that non-genetic mechanisms may also play an important role and information transfer can occur horizontally via a Lamarckian mode of evolution (Álvarez-Arenas et al, 2019). Thus, a population of isogenic cells in the same environment can exhibit single-cell-level stochastic fluctuations in gene expression. Such fluctuations, known as gene expression noise or transcriptional noise, can result in isogenic cells ‘making’ entirely different decisions with regard to their phenotype and hence, their ability to adapt themselves to the same environmental perturbation (Balázsi et al, 2011; Farquhar et al, 2019; Engl, 2019). Transcriptional noise can arise from the intrinsic randomness of underlying biochemical reactions or processes extrinsic to the gene (Swain et al, 2002). Regardless, two main characteristics of gene expression noise are its amplitude and memory. Amplitude, often measured by the coefficient of variation, defines how far cells deviate from the average. Memory describes the time for which cells remain deviant once they depart from the average (Acar et al, 2005; Charlebois et al, 2011). Thus, it follows that the effect noise produces is likely reversible and hence, underscores its importance in phenotypic switching.

Yet another source of noise that can be confounding in phenotypic switching is conformational noise (Mahmoudabadi et al, 2013) that stems from the ‘structural’ plasticity of the IDPs that lack rigid 3D structure and exist as conformational ensembles instead (Wright and Dyson, 2015; Turoverov et al, 2019). Because of their conformational dynamics and flexibility, IDPs can interact with multiple partners and are typically located in ‘hub’ positions in protein interaction networks. The collective effect of conformational noise is an ensemble of protein interaction network configurations, from which the most suitable can be explored in response to perturbations. Moreover, the ubiquitous presence of IDPs as transcriptional factors (Staby et al, 2017; Tsafou et al, 2018), and more generally as hubs (Patil et al, 2010; Hu et al, 2017), underscores their role in propagation of transcriptional noise. As effectors of transcriptional and conformational noise, IDPs rewire protein interaction networks and unmask latent interactions (Mahmoudabadi et al, 2013). Thus, noise-driven activation of latent pathways appears to underlie phenotypic switching events such as drug resistance.

The present data suggest that cisplatin resistance in LUAD can arise stochastically in response to drug treatment. While it is possible that such resistance that is spontaneously, or randomly, acquired during the course of treatment can be due to random genetic mutations or stochastic non-genetic phenotype switching (Pisco et al, 2013), our observations strongly support a non-genetic mechanism. A double negative feedback loop between ITGB4 and miR-1-3p results in bistability, facilitating a reversible phenotypic switch between cisplatin-sensitive and resistant states. A recent study on oxaliplatin chemotherapy in pancreatic ductal adenocarcinoma (Kumar et al, 2019) where the authors studied a coarse-grained stochastic model to quantify phenotypic heterogeneity in a population of cancer cells also mirrors the present findings suggesting the phenomenon may be applicable to many different cancers. Furthermore, a random population of cisplatin-resistant LUAD cells is heterogeneous and comprises individuals that either express high or low levels of ITGB4. These cells therefore are either more resistant or less resistant to cisplatin, respectively. However, when purified to homogeneity (>99%) and plated separately, the purified population recreates the heterogeneity. Taken together, these observations not only corroborate the bistability predicted by the model but also highlight the role of phenotypic switching in generating population heterogeneity in cancer.

Additionally, it is important to note that, the interaction of ITGB4 with the intrinsically disordered PXN is critical since perturbing this interaction renders the cells sensitive. It is important to note that, in addition to the LD domains and the LIM domains, PXN also contains an SH3 domain-binding site and SH2 domain-binding sites (Salgia et al, 1995). Together, these motifs serve as docking sites for cytoskeletal proteins, tyrosine kinases, serine/threonine kinases, GTPase activating proteins, and a host of other adaptor proteins that recruit additional enzymes into complex with PXN. Thus, consistent with the functions of an IDP in a hub position, PXN serves as a docking protein to recruit signaling molecules to the FA complex and thereby, coordinate downstream signaling (Schaller, 2001, Oncogene). It is now well recognized that cellular protein interaction networks are organized as scale-free networks and hence, are remarkably resilient to perturbations (Barabasi and Albert, 1999; Barabasi, 2009). Thus, while disabling minor nodes does not significantly affect the continuity and hence, functionality of the network, attacking the critical nodes can incapacitate the entire network (Schwartz et al, 2002). Thus, it follows that PXN appears to constitute a critical hub and its malfunction accounts for the failure of the cisplatin-resistant cells to tolerate the drug. Additional studies elucidating the structural biology of PXN and the PXN/ITGB4/FAK complex that are currently underway in our laboratories should provide a deeper understanding of the role of these molecules in cisplatin resistance.

Although, IDPs in general have not been much appreciated as therapeutic targets since their inability to adopt well-defined structures provides significant obstacles for developing ligands that regulate their behaviors, emerging evidence indicates that indeed, they can be specifically targeted (Wójcik et al, 2018; Neira et al, 2017; Martin-Yken et al, 2016; Yu et al, 2016; Berg, 2011; Jung et al, 2015; Ambadipudi and Zweckstetter, 2016). In fact, even transcription factors that were never the favourite drug targets, are now emerging as tractable to drug development (Tsafou et al, 2018). Consistent with this growing body of evidence, we have identified a series of FDA-approved drugs including carfilzomib that perturb interactions involving an IDP and may be repurposed for alleviating cisplatin resistance in LUAD. By extrapolation, it is likely that some of these drugs may be effective in several other cancer types in which cisplatin therapy is administered. Thus, carfilzomib and the other drugs we identified in this study can not only hasten clinical trials in future, but may make the availability of the drug, if successful, more cost effective as well. Finally, it is conceivable that using the antibody-drug conjugation technology, carfilzomib or any of the other drugs identified in this study could potentially be conjugated to the ITGB4 antibody and delivered to the tumor site with high specificity (Yao et al, 2016).

Yet another phenomenon that contributes to cisplatin resistance in NSCLC is epithelial to mesenchymal transition (EMT) (Fischer et al, 2015; Zheng et al, 2015). Consistently, we observed that the Smad and TGF-β pathways that are known to be associated with EMT are affected by knocking down ITGB4 and PXN suggesting that there is crosstalk between the molecules that regulate EMT and signaling via the FA complex. Therefore, combining therapies that target the molecules involved in these two processes may prove more effective in treating these patients.

As mentioned before, the role of the FA complex in serving as a conduit through which mechanical force and regulatory signals are transmitted between the extracellular matrix and an interacting cell is well established (Chen et al, 2003). Furthermore, the role of PXN in cisplatin resistance has also been recognized (Wu et al, 2014). However, as far as we are aware, the involvement of ITGB4 and the interaction between these molecules with FAK in modulating cisplatin resistance has not been reported. As summarized in **Fig. 8F**, these results suggest that PXN and ITGB4 are not only involved in tumor invasion and metastasis but are also required for regulating transcription of various genes involved in maintaining genomic stability and mitochondrial function (Fig. 8F). These findings support a new role for the FA complex in cancer, particularly LUAD. Finally, the observation that these FA components may potentially serve as drug response predictors using a blood-based assay, and the identification of FDA-approved drugs that can potentially be repurposed to address drug resistance can have a significant impact on LUAD and perhaps, even other types of cancers that respond to cisplatin.

## Materials and methods

### Cell lines and reagents

Lung cancer cell lines (H23, H358, SW1573, H441, H2009, H522, H1650, H596, H1437, and H1993) were obtained from American Type Culture Collection (ATCC) (Manassas, VA, USA). All cell lines were cultured in RPMI 1640 medium (Corning) supplemented with fetal bovine serum (FBS) (10%), L-glutamine (2 mM), penicillin/streptomycin (50 U/ml), sodium pyruvate (1 mM), and sodium bicarbonate (0.075%) at 37°C, 5% CO_2_. Cisplatin was provided by City of Hope National Medical Center clinics. Anti-integrin beta 4 antibody, clone 8 was purchased from MilliporeSigma (Burlington, MA, USA). FDA-approved drugs were purchased from Selleck Chemicals (Houston, TX, USA).

### Antibodies

Antibodies against ITGB4, FAK, phospho-FAK (Y397), γH2AX, p27, phospho-Rb (S807/811), USP1 were purchased from Cell Signaling Technology (Danvers, MA, USA). Antibodies against ITGA7, ITGA6, PXN, MET, G3BP1, VDAC1, and agarose-conjugated antibodies (ITGB4, FAK, PXN) were purchased from Santa Cruz Biotechnology (Dallas, TX, USA). Cyclin D1 antibody was purchased from Invitrogen (Waltham, MA, USA). Phospho-PXN (Y31) and PARP antibodies were purchased from Abcam (Cambridge, UK). CD63 antibody was purchased from System Biosciences (Palo Alto, CA, USA). β-actin antibody was purchased from Sigma-Aldrich (St. Louis, MO, USA).

### Western blotting

Cell lysates were prepared with 1X RIPA buffer (MilliporeSigma) and denatured in 1X reducing sample buffer at 95°C for 5 min. Protein samples (15 μg) were run on 4-15% TGX gels (Bio-Rad, Hercules, CA, USA) and transferred onto nitrocellulose membranes (Bio-Rad). Blots were blocked with 5% non-fat milk in TBS-T for 1 hour at room temperature and probed with primary antibody diluted in 2.5% BSA in TBS-T overnight at 4°C. After three washes with TBS-T, blots were incubated with HRP-conjugated secondary antibodies for 2 hours at room temperature. After three more washes, bands of interest were visualized via chemiluminescence using WesternBright ECL HRP substrate (Advansta, Menlo Park, CA, USA) and imaged with the ChemiDoc MP imager (Bio-Rad).

### Quantitative real-time PCR

Quantitative real-time PCR (qPCR) reactions were performed using TaqMan Universal PCR Master Mix (Thermo Fisher Scientific, Waltham, MA) and analyzed by the Quant Studio7 Real-timePCR system (Life Technologies, Grand Island, NY). Total RNA isolation and on-column DNase digestion from cells were performed basing on the manufacturer’s protocol RNeasy Plus Mini Kit (Qiagen Cat #: 74134). 1 ug of RNA was used to synthesize the cDNA according to the one step cDNA synthesis kit from QuantaBio (Cat#: 101414-106). TaqMan probes for HS99999905 –GAPDH, HS00236216-ITGB4, HS01104424-PXN, HS00174397-ITGB1, HS00164957-ITGB2, HS01001469-ITGB3, HS01565584-MET, HS04978484-VDAC1, HS00428478-G3BP1 and HS00163427-USP1 were purchased from ThermoFisher (Waltham, MA). The mRNA expression was analyzed using multiplex PCR for the targeted gene of interest and GAPDH as reference using two independent detection dyes FAM probes and VIC probes respectively. Relative mRNA expression was normalized to GAPDH signals and calculated using the ddCt method.

### siRNA Transfection

Knockdown of ITGB4 (Cat #: SR302473C), FAK (Cat #: SR303877C), USP1 (Cat #: SR305052B), and VDAC1 (Cat #: SR305067C) at the mRNA level was executed using siRNAs purchased from OriGene Technologies (Rockville, MD, USA). Knockdown of PXN was achieved by siRNA purchased from Life Technologies Corporation (Cat #: 4392421). JetPRIME transfection reagent (Polyplus Transfection, Illkirch, France) was used to transfect the siRNAs according to the manufacturer’s protocol. Cells were seeded in 6-well plates (200,000 cells/well) and allowed to adhere overnight. Next day, 10 nM siRNA was transfected with 4 μl jetPRIME reagent in complete growth medium for each well. Cell growth medium was changed the next day and expression was detected 72 h post-transfection by immunoblot.

### Cell viability assay

Cell Counting Kit-8 (CCK-8) was purchased from Dojindo Molecular Technologies (Rockville, MD, USA). Cells were seeded on a 96-well plate and allowed to adhere in complete medium for 24 hours. Test compounds were added to 100 µl of medium at the indicated concentrations for 72 hours. Ten μl of the CCK-8 reagent were added to each well and absorbance at 450 nm was measured using a Tecan Spark 10M multimode microplate reader.

### Scratch wound healing assay

Cells were seeded on a 96-well ImageLock (Essen BioScience, Ann Arbor, MI, USA) plate to reach 90% confluence by the next day. After cell adherence, 96 uniform wounds were created simultaneously using the WoundMaker (Essen BioScience) tool. Cells were washed once with serum-free medium and replenished with complete medium. To monitor wound healing, the plate was placed in the IncuCyte S3 Live-Cell Analysis System (Essen BioScience) and images were acquired every hour. Data analysis was generated by the IncuCyte software using a set confluence mask to measure relative wound density over time.

### Cell proliferation and apoptosis assay

Cell proliferation assays were performed using cell lines stably transfected with NucLight Red Lentivirus (Essen Bioscience) to accurately visualize and count the nucleus of a single cell. Cells were seeded on a 96-well plate and allowed to adhere for 24 h. Test compounds were added at indicated concentrations. Caspase-3/7 Green Apoptosis Reagent (Essen Bioscience) was also added as a green fluorescent indicator of caspase-3/7-mediated apoptotic activity. To monitor cell proliferation and apoptosis over time, the plate was placed in the IncuCyte S3 Live-Cell Analysis System (Essen BioScience) and images were acquired every 2 hours. Data analysis was generated by the IncuCyte software using a red fluorescence mask to accurately count each cell nucleus and a green fluorescence mask to measure apoptosis over time.

### Immunoprecipitation (IP)

Cells were lysed in the Pierce™ IP Lysis Buffer purchased from Thermo Fisher Scientific and 1 mg of protein was allowed to bind overnight in 4°C to agarose-conjugated antibodies (Santa Cruz Biotechnology): ITGB4 (Cat #: sc-13543 AC), FAK (Cat #: sc-271195 AC), PXN (Cat #: sc-365379 AC). IP beads were washed 5 times with 1X RIPA buffer and denatured in 2X reducing sample buffer at 95°C for 5 min. Western blots according to aforementioned protocol were performed to determine IP results.

### 3D spheroid assay

3D spheroid experiments were performed using cell lines stably transfected with NucLight Red Lentivirus (Essen Bioscience) to visualize red fluorescence as an indicator of cell viability. Cells were seeded on a 96-well ultra-low attachment plate and allowed to form spheroids overnight. Drug treatment was added as indicated along with Cytotox Green Reagent (Essen BioScience), used as a green fluorescence indicator of cell death due to loss of cell membrane integrity. To monitor cell proliferation and apoptosis over time, the plate was placed in the IncuCyte S3 Live-Cell Analysis System (Essen BioScience) and images were acquired every 2 hours. Data analysis was generated by the IncuCyte software using a red fluorescence mask to accurately measure intensity and area of red fluorescence, indicating spheroid viability and a green fluorescence mask, indicating cell death.

### Cell cycle analysis

H2009 cells were harvested and pelleted after 72 h following siRNA transfection. Ice cold 70% ethanol was added to the pellet with mild vortexing to fix the cells. The fixed cells were kept at 4°C for PI staining. FxCycle™ PI/RNase Staining solution from Invitrogen was used for staining the DNA according to the manufacturer’s protocol prior the FACS analysis. Univariate model of Watson (Pragmatic) was used for cell cycle analysis.

### Confocal microscopy

3D spheroids were seeded and imaged in 96-well clear ultra-low attachment microplates (Corning) using Zeiss LSM 880 confocal microscope with Airyscan at the Light Microscopy/Digital Imaging Core Facility at City of Hope. Images were processed using ZEN software and analyzed using ImageJ (Schneider, C. A.; Rasband, W. S. & Eliceiri, K. W. (2012), “NIH Image to ImageJ: 25 years of image analysis”, Nature methods 9(7): 671-675, PMID 22930834).

### Seahorse XF Cell Mito Stress Test metabolic assay

Cells were seeded in complete growth medium on a Seahorse XF Cell Culture Microplate (Agilent Technologies, Santa Clara, CA, USA) to reach 90% monolayer confluence by the next day. One day prior to assay, 5 μM cisplatin was added for 24 h. On the day of the assay, mitochondrial inhibitor compounds were added to injection ports of the XFe96 FluxPak sensor cartridge at a final concentration of: oligomycin 1 μM, FCCP 1 μM, rotenone/antimycin A 1 μM each. Culture medium was changed to assay medium: Seahorse XF RPMI medium supplemented with 1 mM sodium pyruvate, 2 mM L-glutamine, and 10 mM glucose. After completion of assay, cells were immediately stained with Hoechst dye and imaged using BioTek. Images were analyzed with QuPath (Bankhead, P. et al. (2017). QuPath: Open source software for digital pathology image analysis. Scientific Reports. https://doi.org/10.1038/s41598-017-17204-5) to obtain number of cells in each well and normalize data according to cell number.

### ROS production assa

Cells were seeded in a 96-well plate and placed in an incubator at 37°C for 72 h. 50 μl of medium from each well was transferred to another 96-well plate to measure ROS production with ROS-Glo™ H_2_O_2_ Assay (Promega, Madison, WI, USA). Remaining plate with cells was used to perform CellTiter-Glo® Luminescent Cell Viability Assay (Promega) to normalize ROS data to number of viable cells. Luminescence was measured using a Tecan Spark 10M multimode microplate reader.

### γH2AX foci staining and analysis

Cells were seeded (50,000 cells/well) on glass cover slips coated with 0.1% gelatin (Millipore) in a 12-well plate. Next day, 5 μM cisplatin was added for 24 h. Cells were fixed in 4% formaldehyde for 30 min at room temperature and blocked. Primary antibody against γH2AX (Cell Signaling Technology) was incubated in 4°C overnight. Then secondary antibody was incubated for 2 hours at room temperature. Cover slips were mounted on glass slides and imaged with Zeiss LSM 880 confocal microscope at the Light Microscopy/Digital Imaging Core Facility at City of Hope. Using QuPath (Bankhead, P. et al. (2017). QuPath: Open source software for digital pathology image analysis. Scientific Reports. https://doi.org/10.1038/s41598-017-17204-5), green fluorescent subcellular particles were counted in each nucleus to obtain γH2AX foci count per cell.

### Exosome isolation and analysis

200 μl of serum from patient or healthy donor was taken and diluted with PBS to 5ml. The diluted serum was filtered through 0.22-micron syringe filter and ultra-centrifuged at 90,000 × g for 90 min. The supernatant was decanted, and the pellet was washed and centrifuged in 15 ml of PBS as before. The PBS was decanted, and pellet was suspended in 50 μl of 1X RIPA buffer containing protease inhibitors for 30 min on ice and transferred to 1.5 ml tube for quantification and denaturation as mentioned above

### Chromatin Immunoprecipitation

Briefly, five million formaldehyde-fixed cells were lysed in 200 μl of SDS lysis buffer and diluted to 2 ml in ChIP dilution buffer in the presence of protease inhibitors. Lysates were sonicated using Bioruptor PICO for 3 cycles and each cycle has 10 repeats of 30 sec pulse and 30 sec break. Lysates were precleared in salmon sperm DNA and protein A agarose by centrifugation. Prior to addition of antibody, 10% of the lysate was used for input and the remaining lysate was divided into two equal parts, one for IgG control and other for H3K27 acetylated antibody from Diagenode. Downstream processing of the chromatin bound antibody was done as per the manufacturer’s protocol for EZ-magna ChIP A/G (Millipore, Temecula, CA). The extracted DNA was used for SYBR green based qPCR assay using the primers sequences Upstream USP1R-5’-AGGTTCACAGCATTCTCAATCC-3’, Upstream USP1F-CAGTGCCTGTGAAACTTTGGA, Promoter USP1F-CTCAGCTCTACAGCATTCGC and Promoter USP1R-GGCCATCCAATGAGACAAGG. The data was analyzed based on the percentage of input.

### In-silico prediction of paxillin conformational ensemble

Using in-silico modeling and enhanced sampling MD simulations, we predicted the conformational ensemble of Paxillin N-terminal domain between LD2 and LD4 bound to FAK. Starting from the crystal structures of FAK bound to the LD2 and LD4 motifs of Paxillin (pdb IDs 1OW8 and 1OW7 [2]), we modeled the disordered region of Paxillin (∼110 residues) between LD2 and LD4 as a random coil using Modeller. The N and the C termini of Paxillin and FAK were capped by the acetyl and N-methyl acetamide groups respectively. This structure was subjected to temperature replica exchange simulations (REMD) [5] using the AMBER ff14SBonlysc force-field and the implicit solvation model GBneck2, as recommended in the AMBER16 manual. Twenty-six replicas between 280K and 480K were used and the individual temperatures were chosen to maintain an exchange success rate of 20%. To maintain structural integrity when subjected to high temperature during REMD, the backbone atoms of the helical regions of FAK and the bound LD2 and LD4 domains of Paxillin were position-restrained with a force of 5 kcal/mol. During REMD, the temperature was maintained using the Langevin thermostat with a collision frequency of 1.0. To accelerate the MD, the system was subjected to hydrogen mass repartitioning, which allowed a 4 femtosecond timestep to be used. The replicas were initially minimized using the conjugate gradient method followed by gradual heating to their respective temperatures over 1 ns. The REMD simulation was carried out for 1.15 μs for each replica, followed by clustering of the lowest temperature conformations by the Paxillin Cα atoms using hierarchical agglomerative clustering. The representative structures from the top 3 most populated clusters were used for virtual screening.

### Virtual ligand screening to identify small molecule inhibitors of Paxillin-FAK interaction

The top Paxillin conformations obtained from REMD were scanned for druggable pockets using the program FindBindSite (FBS). The top ranked pockets in each Paxillin conformation were further screened by proximity to the FAK interface. In total, 4 pockets among 3 Paxillin conformations were selected for virtual screening. The protein structures were prepared using Maestro (Schrodinger™, LLC). The SelleckChem FDA-approved drug library containing approximately 1400 compounds was then docked to each pocket separately using Glide standard precision. The ligand library was prepared using the LigPrep module of Maestro and all possible protonation states at neutral pH were generated. During docking, the protein atoms were scaled by 0.8 and 10 docked poses per ligand were retained. Next, the best docked pose for each ligand was selected by Glide score and was optimized by reassigning the side chains within 5Å of the docked ligand using Prime followed by minimization of the entire complex using MacroModel (Schrodinger™, LLC). Using MacroModel, a crude binding energy score was generated for each docked complex by subtracting the sum of individually solvated protein and ligand energies from the energy of the solvated complex. This score was used to select the top 50 ligands from each binding pocket, which were then subjected to thorough optimization and binding free energy calculation using the MMGBSA method in PrimeX. The top 10 compounds by binding free energy from each binding pocket were selected for experimental testing.

### Fluorescence-activated cell sorting (FACS) and analysis

Cells were trypsinized and resuspended (5 million) in PBS with 2% FBS. Cells were stained with ITGB4 antibody conjugated to Alexa Fluor® 488 (5 μl/1 million cells) (R&D Systems, Minneapolis, MN, USA) and Propidium Iodide Ready Flow™ Reagent (1 drop/1 million cells) (Invitrogen) for 30 min at 4°C. The Analytical Cytometry Core Facility at City of Hope carried out and assisted all FACS sorting and analysis experiments. Gates were set to sort cell populations having low 10% and high 10% expression of ITGB4 using the FACSAria™ Fusion (BD Biosciences, San Jose, CA, USA). Sorted cells were immediately cultured in 12-well plates and treated with cisplatin (1 μM) for 48 h. Then, equal number of untreated and treated cells were collected and stained with same reagents as above. FACS analysis was performed to determine shifts in cell population using the Attune NxT Flow Cytometer (Invitrogen).

### Mathematical modeling

Bifurcation diagram was obtained MATCONT (cite: https://dl.acm.org/citation.cfm?id=779362). Next, Random circuit perturbation (RACIPE) algorithm was run on the two-node network – ITGB4/ miR-1-3p. The continuous gene expression levels were obtained as output with randomly chosen parameters for the regulatory links. The algorithm was used to generate 100,000 mathematical models, each with a different set of parameters for the following ODEs:

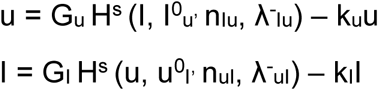

where, u denotes miR-1-3p and I denotes ITGB4. G_u_ and G_I_ are the maximum production rates of miR-1-3p and ITGB4 respectively. And, k_u_ and k_I_ are their innate degradation rates respectively.

## Acknowledgements

MKJ was supported by Ramanujan Fellowship (SB/S2/RJN-049/2018) provided by SERB, DST, Government of India

## Supplementary information

Equations:

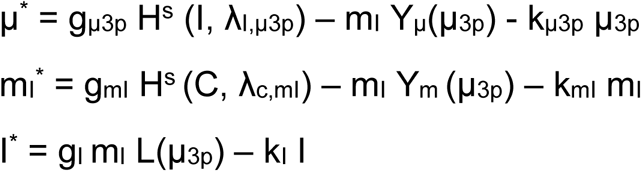

where H^S^ is the shifted Hill function, defined as H^S^ (B,λ) = H^-^(B)+λH^+^ (B), H^−^ (B) =1/ [1+(B / B_0_)^nB^], H^+^ (B) =1−H^−^ (B) and λ is the fold change from the basal synthesis rate due to protein B. λ >1 for activators, while λ <1 for inhibitors.

The total translation rate:

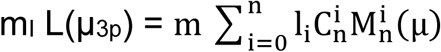

The total mRNA active degradation rate:

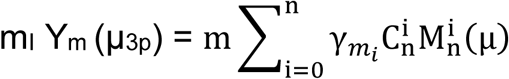

The total miR active degradation rate is

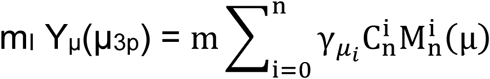

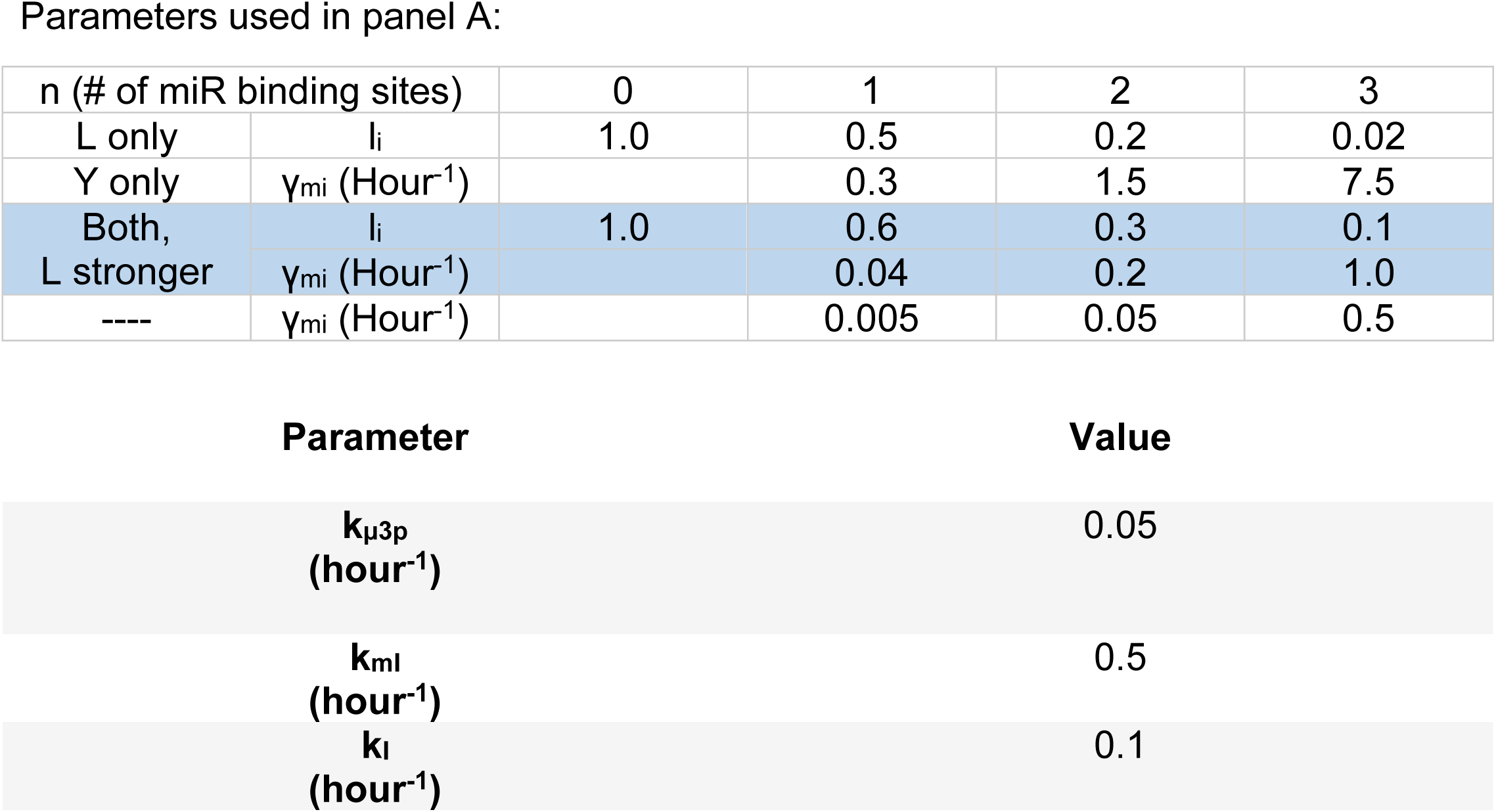
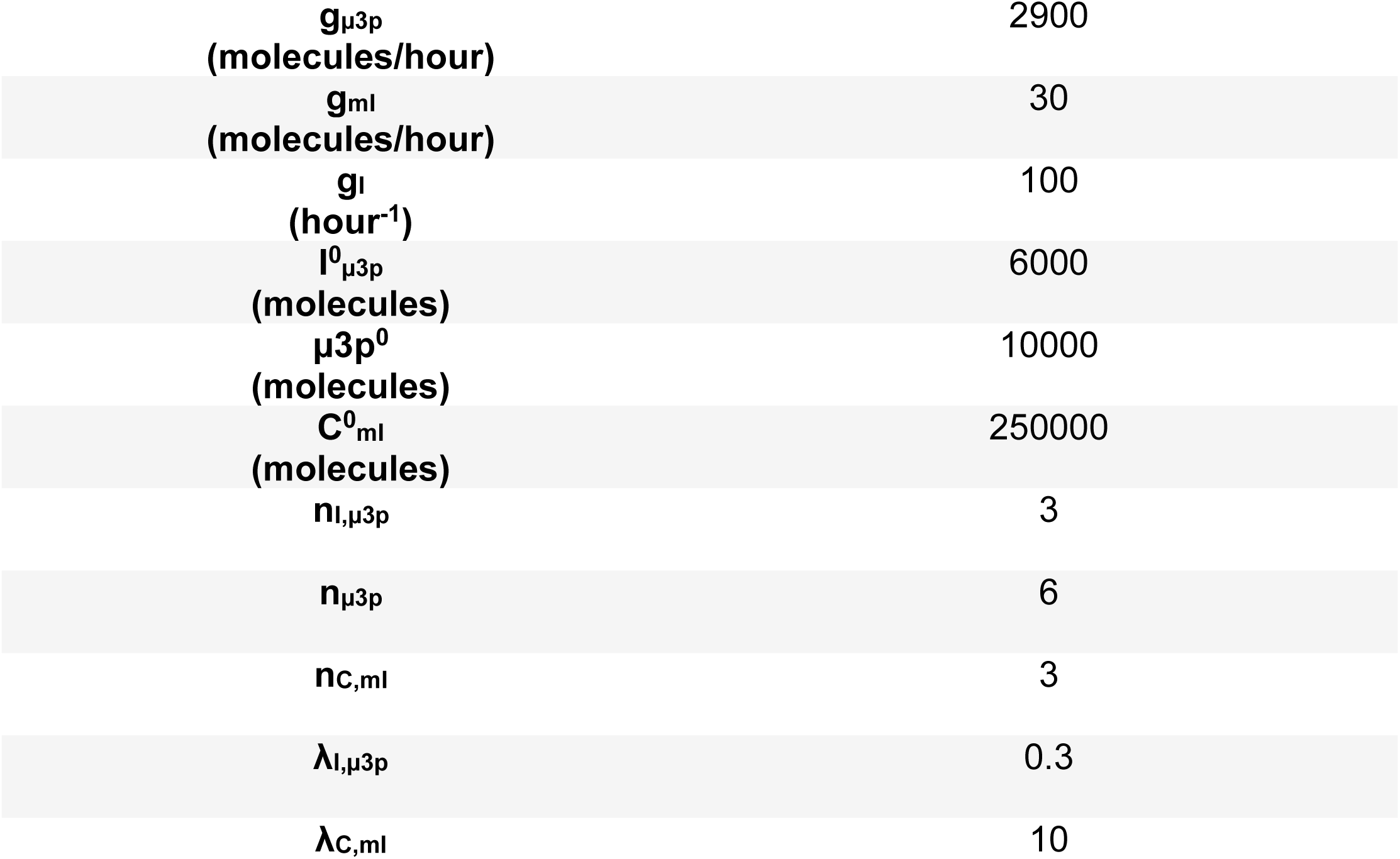

The feedback loop was constructed based on data reported in the manuscript (ITGB4 inhibits miR-1-3p) and publicly available data (miR-1-3p inhibits ITGB4) - https://www.genecards.org/cgi-bin/carddisp.pl?gene=ITGB4.

The parameters for microRNA-mediated dynamics were estimated from our previous models for microRNA-mediated regulation of EMT (Lu et al. 2013).

## Supplementary Figure Legends

**Supplementary Figure 1.**
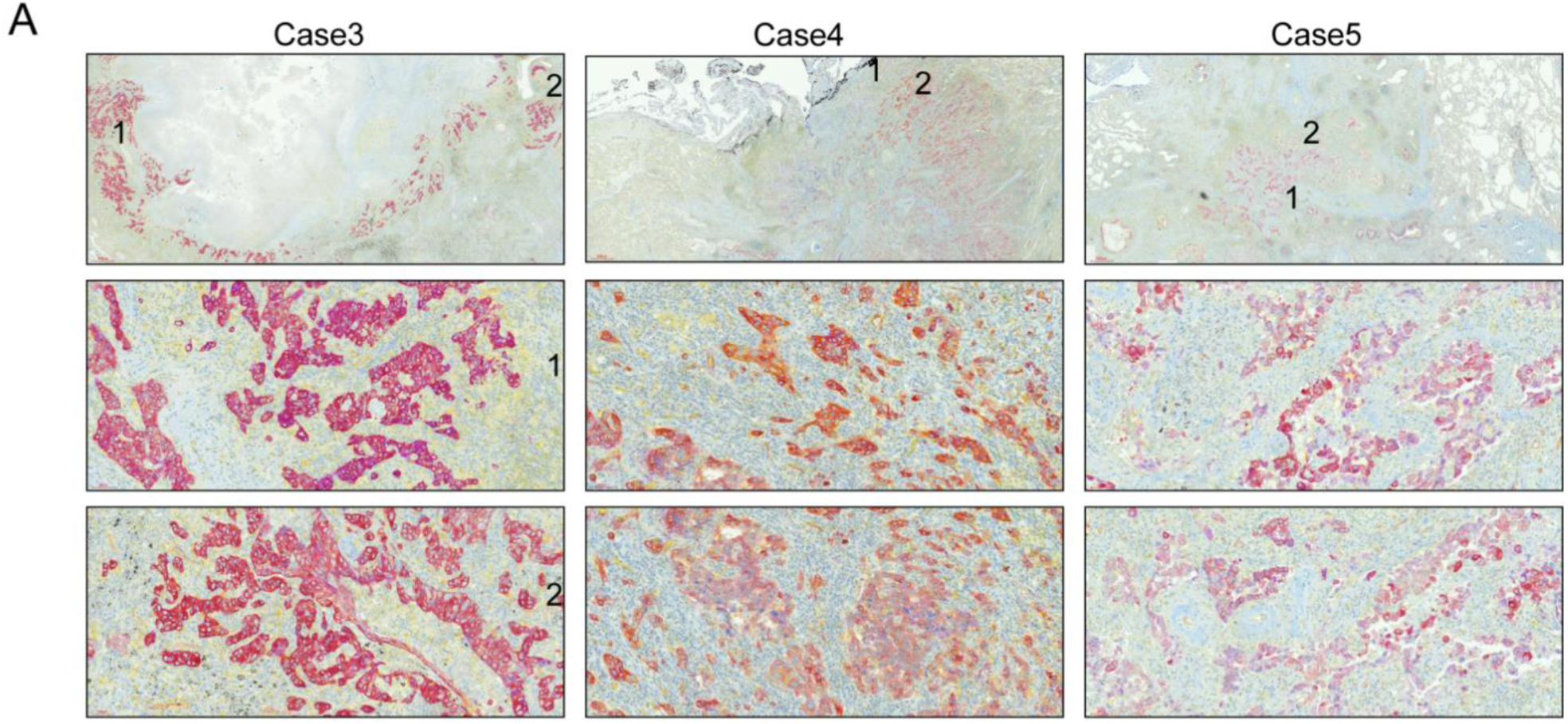
Variability of expression of PXN and ITGB4 in tumor tissue samples. **A)** Immunohistochemistry staining of lung adenocarcinoma tumor tissue showed that Case 1 had high PXN and intermediate ITGB4 expression. In Case 2, PXN expression is low and ITGB4 expression is high. In Case 3, both PXN and ITGB4 expression are low.

**Supplementary Figure 2.**
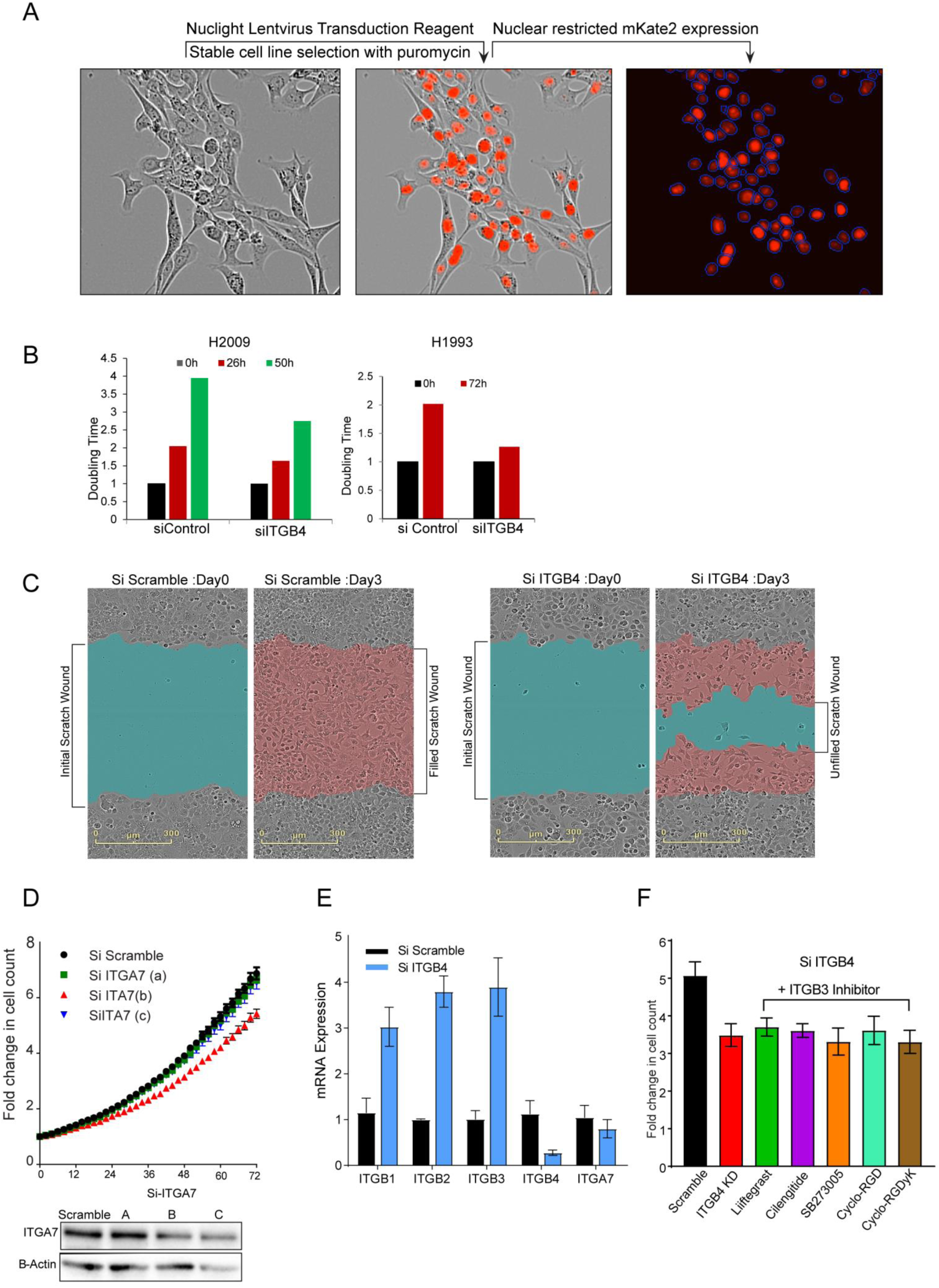
Cell proliferation, apoptosis, and wound healing assay methods. **A)** Stable cell lines were generated to express nuclear mKate2 red fluorescent protein (RFP) using the NucLight Red Lentivirus Reagent (Essen BioScience). Upon selection with puromycin, cells were analyzed with the IncuCyte software to create a mask around each individual nucleus and obtain accurate real-time cell counts for proliferation assays. **B)** Doubling times for H2009 and H1993 cells transfected with scramble siRNA (siControl) were measured and compared to that of cells with ITGB4 knockdown (siITGB4). **C)** Scratch wound healing assays were performed by creating an initial scratch wound with the WoundMaker tool. Wound closure was quantitated by monitoring cells that migrated to fill the initial wound. Cells transfected with scramble siRNA (Si Scramble) were able to completely fill the scratch wound by day 3 whereas ITGB4 knockdown cells (Si ITGB4) were not. **D)** H2009 cells transfected with10 nM of siRNA constructs A, B, and C, targeting ITGA7 had minimal effect on proliferation. **E)** Knocking down ITGB4 increased mRNA expression of other integrin beta forms such as ITGB1, ITGB2, and ITGB3 but had no significant effect on the expression of ITGA7. **F)** In order to nullify the effect of ITGB3 rescue, H2009 cells transfected with siRNA ITGB4 were treated with ITGB3 inhibitors. There was no significant effect in the fold change in proliferation compared to the ITGB4 knockdown cells

**Supplementary Figure 3.**
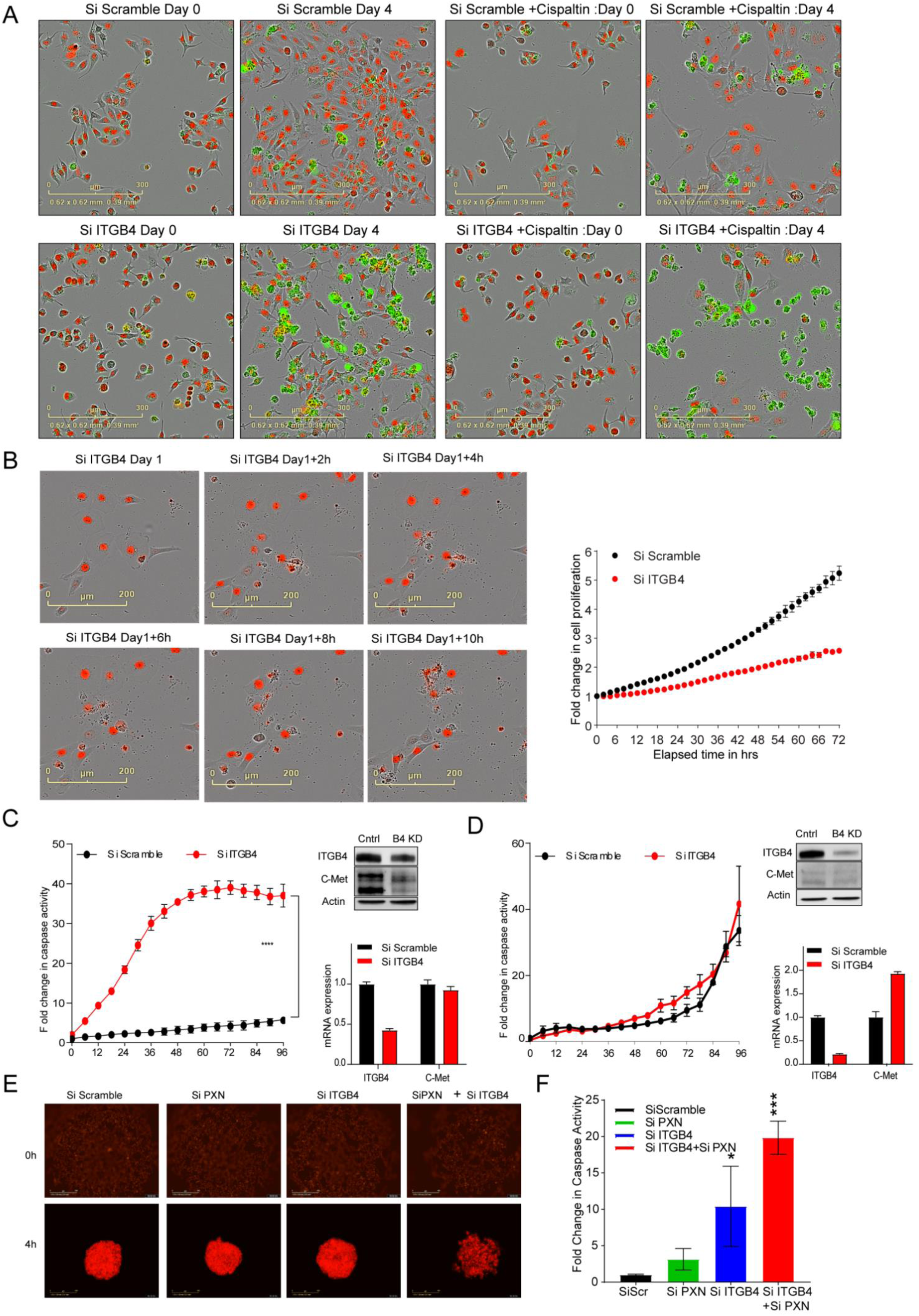
ITGB4 knockdown alone induces varying effects in different LUAD cell lines. **A)** Proliferation and apoptosis assays were executed with stable cell lines expressing nuclear RFP and the IncuCyte Caspase-3/7 Green Reagent (Essen BioScience), which emits green fluorescence when cleaved by activated caspase-3/7. Apoptosis is induced with knockdown of ITGB4 (Si ITGB4) by day 4 and enhanced with added cisplatin (Si ITGB4+Cisplatin). **B)** Nuclear RFP-expressing cell line H1650 was transfected with ITGB4 siRNA (Si ITGB4) and monitored in real-time with the IncuCyte. Over the course of 10 h, ITGB4 knockdown induced cells with an intact membrane to undergo anoikis-like bursting, attenuating cell proliferation in Si ITGB4 (red) cells compared to control (black). **C)** H1993 cells harbor MET amplification and upon ITGB4 knockdown, MET expression at only the protein level decreases but no change in the mRNA expression, indicating ITGB4 is required for MET protein stability but not for transcriptional regulation. However, ITGB4 knockdown increased apoptosis significantly compared to control cells. **D)** H2009 cells that do not have amplified MET did not show any significant change in the protein as well as mRNA expression. They also did not show significant increase in apoptosis with ITGB4 knockdown. **E)** H2009 cells expressing nuclear RFP were seeded in an ultra-low attachment 96-well plate to facilitate spheroid formation. After 4 h, double knockdown of PXN and ITGB4 impeded cells from forming a compact spheroid as observed in control and single knockdown conditions. **F)** In H2009 cells, single knockdown of ITGB4 (Si ITGB4) significantly induced apoptosis and double knockdown of PXN and ITGB4 (Si ITGB4+Si PXN) had an enhanced effect.

**Supplementary Figure 4.**
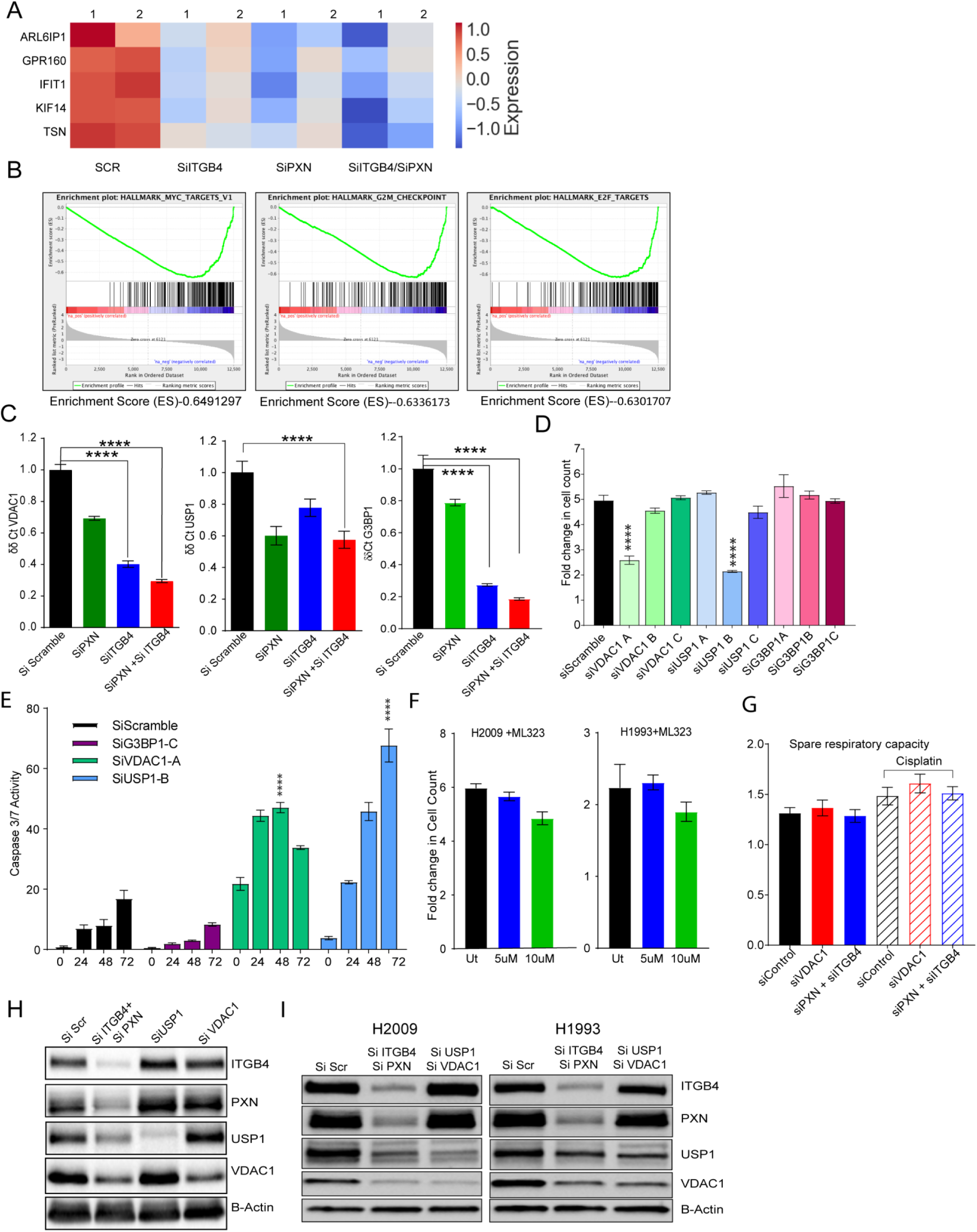
Both ITGB4 and PXN knockdown downregulate genes involved in a common pathway. **A)** Total RNA sequencing after single and double knockdown of PXN and ITGB4 revealed that 5 common genes were downregulated compared to control cells (SCR). **B)** Major pathways affected by double knockdown of PXN and ITGB4 were MYC targets, G2-M checkpoint, and E2-F targets. **C)** qPCR results with single and double knockdown of PXN and ITGB4 confirmed downregulation of VDAC1, USP1, and G3BP1 at the mRNA level. **D)** SiRNA constructs A, B, or C against top four genes (KIF14, VDAC1, USP1, and G3BP1) down regulated were tested to determine the construct with maximum effect on inhibiting proliferation. KIF14 construct C, VDAC1 construct A, and USP1 construct B showed significant effect on cell proliferation. **E)** The selected constructs were tested for induction of caspase-3/7 activity by tracking green fluorescence using live cell imaging and analysis for 72 hrs. **F)** H2009 and H1993 cells treated with ML323, a USP1 inhibitor, did not undergo significant changes in proliferation compared to untreated cells (black). **G)** Spare respiratory capacity, which is the ratio of maximal respiration vs. basal respiration, did not show any significant change between control and knockdown cells. **H)** Double knockdown of PXN and ITGB4 (Si ITGB4+Si PXN) decreased expression of USP1 and VDAC1. However, knocking down USP1 (Si USP1) or VDAC1 (Si VDAC1) did not affect expression of ITGB4 or PXN. **I)** H2009 and H1993 cells with PXN/ITGB4 or USP1/VDAC1 double knockdown confirmed by immunoblot to be used for ROS production assay.

**Supplementary Figure 5.**
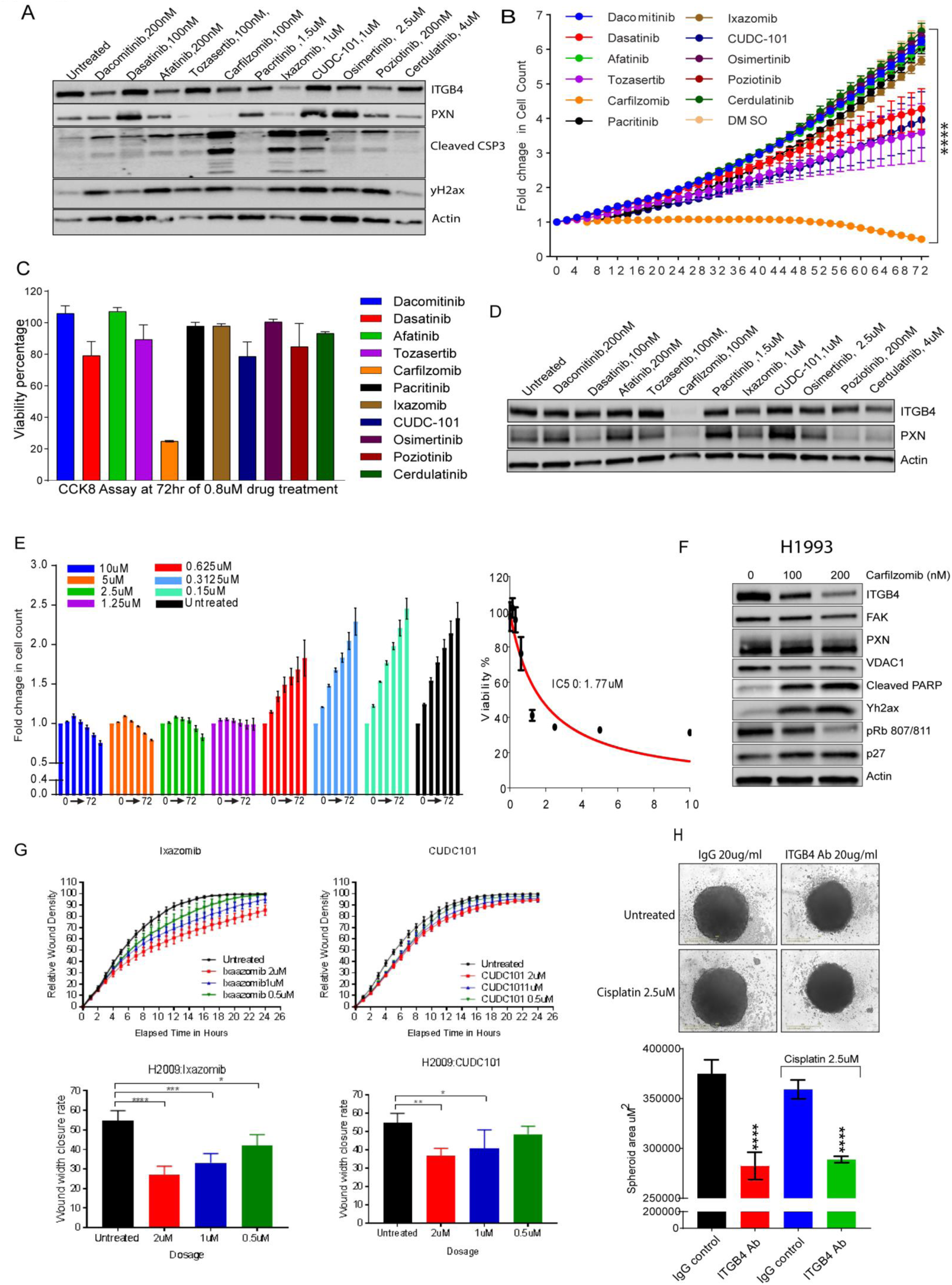
FDA-approved compounds were screened and tested to determine the most effective compound in cisplatin-resistant cell lines. **A)** Eleven FDA-approved compounds that exhibited the greatest inhibitory effect on H358 cells were further analyzed with immunoblot. H358 cells were treated with each compound at the indicated sublethal dose for 72 h. **B)** and **C)** H2009 cells treated with the eleven FDA-approved compounds at 0.08 μM. Cell proliferation over the course of 72 h was monitored and viability percentage compared to untreated was measured using CCK-8 assay. **D)** Immunoblot of H2009 cells treated with sublethal dose of eleven drugs for 72 h showed reduction in both ITGB4 and PXN expression with only carfilzomib treatment. **E)** H1993 cells were treated with an increasing dose of carfilzomib to determine cell proliferation effect with change in dose and determine the IC50. **F)** Immunoblot of H1993 cells treated with carfilzomib showed reduced expression of ITGB4 and phospho-Rb (S807/811) and increased expression of p27, γH2AX, and cleaved PARP, indicating cell cycle inhibition and cell death. **G)** Ixazomib and CUDC-101, two compounds that induced caspase activity, were further analyzed with scratch wound healing assays in H2009 cells by taking into parameters relative wound density and wound closure rate. Only ixazomib impeded cell migration and CUDC-101 did not. **H)** Treating the patient-derived organoids Lcsc4 with 20 μg/ml of ITGB4 antibody decreased the spheroid area significantly, but combination with cisplatin 2.5 μM had no additive effect.

**Supplementary Figure 6.**
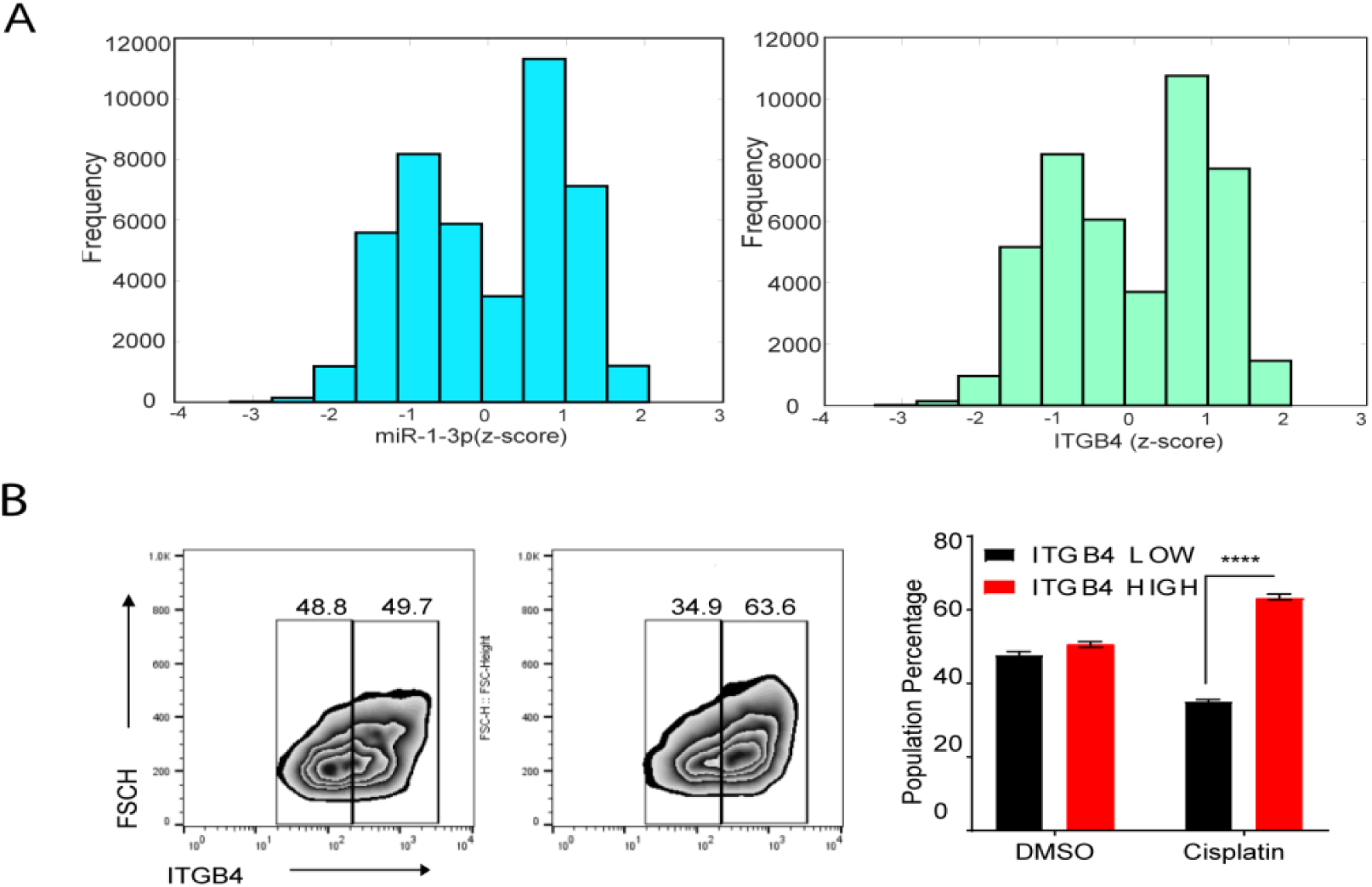
**A)** RACIPE ensemble results (n=100,000) show ITGB4 and miR-1-3p exhibit bimodality: two distinct subpopulations of cells, as shown via z-socre distributions of ITGB4 and miR-1-3p. **B)** FACS analysis using an anti-ITGB4 antibody showed enrichment of high ITGB4 population upon cisplatin treatment in H2009 cells.

**Supplementary Video 1,2**: Wound healing assay, post Scramble or ITGB4 SiRNA transfection in H1993 cells respectively.

**Supplementary Video 3,4,5,6:** Real time increase of green fluorescence indicating activation of caspase 3/7 activity after scramble Si RNA transfection alone or with cisplatin treatment or ITGB4 Si RNA transfection alone or with cisplatin treatment respectively.

**Supplementary Video 7,8:** Cell bursting after scramble knockdown or ITGB4 knockdown in H1650 cell lines respectively.

**Supplementary Video 9,10,11,12:** Live video showing change in H2009 cell line derived spheroid after scramble knockdown or PXN knockdown or ITGB4 knockdown or both PXN and ITGB4 knockdown respectively.

**Table 1.** Genes upregulated in cisplatin-resistant lung adenocarcinoma cell line from Molecular signature database, Broad Institute.

| Original Member | Entrez GeneId | Gene Symbol | GeneDescription |
| --- | --- | --- | --- |
| AF010316 | 9536 | PTGES | Prostaglandin E Synthase |
| AF019770 | 9518 | GDF15 | Growth Differentiation Factor 15 |
| J04164 | 8519 | IFITM1 | Interferon Induced Transmembrane Protein 1 |
| M29366 | 2065 | ERBB3 | V-erb-b2 Erythroblastic Leukemia Viral Oncogenesis |
| M29870 | 5879 | RAC1 | Ras-related C3 Botulinum Toxin Substrate 1 |
| M33882 | 4599 | MX1 | Myxovirus (Influenza Virus) Resistance 1 |
| S80437 | 2194 | FASN | Fatty Acid Synthase |
| U09579 | 1026 | CDKN1A | Cyclin-dependent Kinase Inhibitor 1A |
| U14588 | 5829 | PXN | Paxillin |
| X53587 | 3691 | ITGB4 | Integrin, Beta 4 |
| X74295 | 3679 | ITGA7 | Integrin, Alpha 7 |

**Table 2.** Cox proportional hazards model: Survival ∼ Sex + Stage + Age + PXN * ITGB4.

|  | Estimate | Standard Error | Relative Risk | P-Value |
| --- | --- | --- | --- | --- |
| gender[1] | - 0.04918 | 0.154775 | 0.952008 | 0.750665 |
| stage[S2] | 0.738091 | 0.194204 | 2.09194 | 0.000144 |
| stage[S3] | 1.17346 | 0.192492 | 3.23316 | 1.09E- 09 |
| stage[S4] | 1.31524 | 0.284479 | 3.72563 | 3.78E- 06 |
| age | 2.36E- 05 | 2.14E- 05 | 1.00002 | 0.271914 |
| ENSG00000089159*<br>ENSG00000132470 | 0.000156 | 3.03E- 05 | 1.00016 | 2.47E- 07 |

**Table 3.**
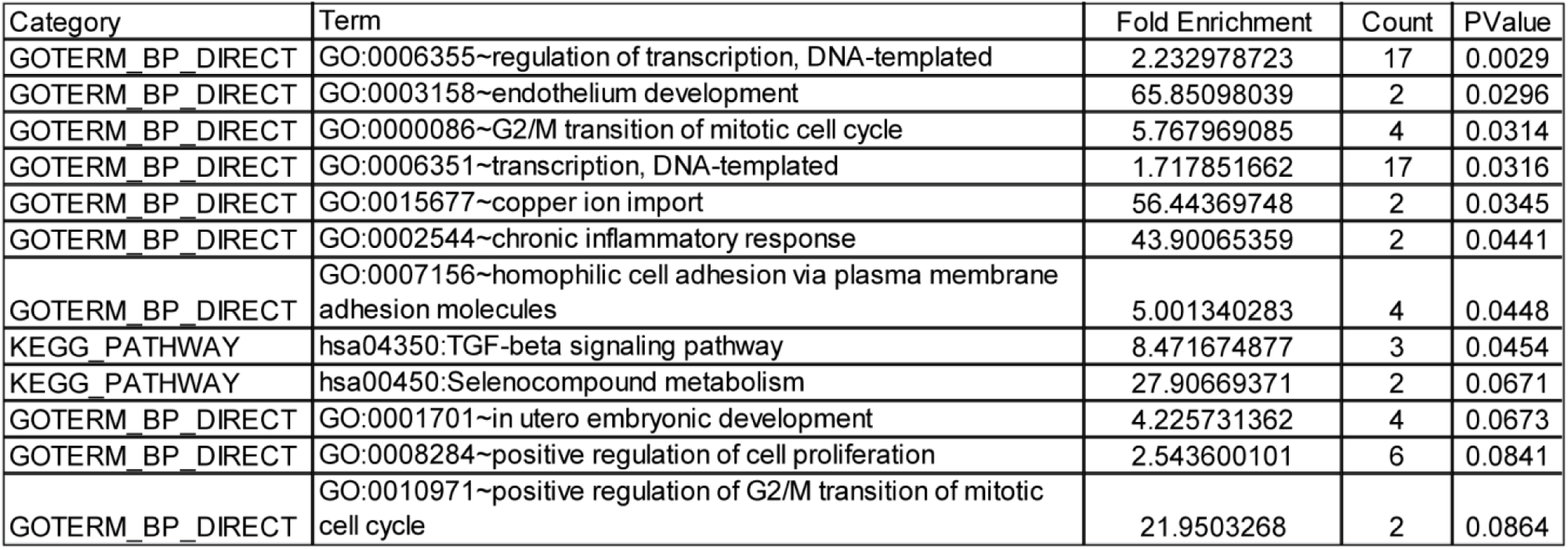
DAVID analysis of 96 unique genes downregulated in double knockdown of PXN and ITGB4.

**Table 4.** DAVID analysis of 206 common genes downregulated in single and double knockdown of PXN and ITGB4.

| Category | Term | Fold Enrichment | Count | PValue |
| --- | --- | --- | --- | --- |
| GOTERM_BP_DIRECT | GO:0006334~nucleosome assembly | 5.759560967 | 8 | 4.79E-04 |
| GOTERM_BP_DIRECT | GO:0007265~Ras protein signal transduction | 6.119533528 | 5 | 0.00891 |
| KEGG_PATHWAY | hsa04810:Regulation of actin cytoskeleton | 3.04717608 | 8 | 0.01475 |
| GOTERM_BP_DIRECT | GO:0001525~angiogenesis | 3.073487691 | 8 | 0.01553 |
| KEGG_PATHWAY | hsa05034:Alcoholism | 3.163381947 | 7 | 0.02182 |
| GOTERM_BP_DIRECT | GO:0017085~response to insecticide | 85.67346939 | 2 | 0.02309 |
| KEGG_PATHWAY | hsa05322:Systemic lupus erythematosus | 3.5815689 | 6 | 0.02467 |
| GOTERM_BP_DIRECT | GO:0032091~negative regulation of protein binding | 6.01217329 | 4 | 0.02854 |
| KEGG_PATHWAY | hsa04070:Phosphatidylinositol signaling system | 4.081039393 | 5 | 0.03263 |
| GOTERM_BP_DIRECT | GO:0060978~angiogenesis involved in coronary vascular morphogenesis | 57.11564626 | 2 | 0.03444 |
| GOTERM_BP_DIRECT | GO:0006203~dGTP catabolic process | 57.11564626 | 2 | 0.03444 |
| GOTERM_BP_DIRECT | GO:0045815~positive regulation of gene expression, epigenetic | 5.527320606 | 4 | 0.03538 |
| GOTERM_BP_DIRECT | GO:0009611~response to wounding | 5.439585358 | 4 | 0.03684 |
| GOTERM_BP_DIRECT | GO:0043928~exonucleolytic nuclear-transcribed mRNA catabolic process involved in deadenylation-dependent decay | 8.862772695 | 3 | 0.04437 |
| GOTERM_BP_DIRECT | GO:0035404~histone-serine phosphorylation | 42.83673469 | 2 | 0.04565 |
| GOTERM_BP_DIRECT | GO:0007052~mitotic spindle organization | 8.567346939 | 3 | 0.04718 |
| GOTERM_BP_DIRECT | GO:0030855~epithelial cell differentiation | 4.895626822 | 4 | 0.04791 |
| GOTERM_BP_DIRECT | GO:0006335~DNA replication-dependent nucleosome assembly | 8.031887755 | 3 | 0.053 |
| GOTERM_BP_DIRECT | GO:0061551~trigeminal ganglion development | 34.26938776 | 2 | 0.05674 |
| KEGG_PATHWAY | hsa04152:AMPK signaling pathway | 3.251559841 | 5 | 0.06526 |
| GOTERM_BP_DIRECT | GO:0042060~wound healing | 4.283673469 | 4 | 0.06621 |
| GOTERM_BP_DIRECT | GO:0032958~inositol phosphate biosynthetic process | 28.55782313 | 2 | 0.06769 |
| GOTERM_BP_DIRECT | GO:0071455~cellular response to hyperoxia | 24.47813411 | 2 | 0.07852 |
| GOTERM_BP_DIRECT | GO:0051290~protein heterotetramerization | 6.119533528 | 3 | 0.08541 |
| GOTERM_BP_DIRECT | GO:0007160~cell-matrix adhesion | 3.807709751 | 4 | 0.08722 |
| GOTERM_BP_DIRECT | GO:0051597~response to methylmercury | 21.41836735 | 2 | 0.08923 |
| GOTERM_BP_DIRECT | GO:0030514~negative regulation of BMP signaling pathway | 5.711564626 | 3 | 0.09605 |
| GOTERM_BP_DIRECT | GO:0060627~regulation of vesicle-mediated transport | 19.03854875 | 2 | 0.09981 |
| GOTERM_BP_DIRECT | GO:0030903~notochord development | 19.03854875 | 2 | 0.09981 |

**Table 5.** IC50 values of H358 cells treated with 11 FDA approved compounds.

| Drug | IC50 (uM) |
| --- | --- |
| Carfilzomib | 0.0008 |
| Tozasertib | 0.0009 |
| Dasatinib | 0.001 |
| Afatinib | 0.009 |
| Dacomitinib | 0.06 |
| Poziotinib | 0.06 |
| Pacritinib | 0.1714 |
| Ixazomib | 0.303 |
| Osimertinib | 0.358 |
| CUDC-101 | 0.501 |
| Cerdulatinib | 1.54 |

**Table 6.** MiRNAs differentially regulated by single and double knockdown of PXN and ITGB4

|  | Activation Score | Predicted Activation | P -value |
| --- | --- | --- | --- |
| ITGB4 Knockdown |  |  |  |
| miR -223 | 2.236 | Activated | 0.0985 |
| miR -210 | 2.449 | Activated | 0.00772 |
| miR-124-3p | 2.077 | Activated | 0.00352 |
| miR-1-3p | 3.128 | Activated | 0.0000133 |
| PXN KD |  |  |  |
| miR -199a-5p | -2.43 | Inhibited | 0.000731 |
| PXN/ITGB4 Knockdown |  |  |  |
| miR-291a-3p | 2.562 | Activated | 1.25E -08 |
| miR -124-3p | 2.292 | Activated | 0.0000371 |
| miR -30c-5P | 2.345 | Activated | 0.0081 |
| miR -1-3p | 2.543 | Activated | 0.00296 |
| miR -199a- 5p | -2.135 | Inhibited | 0.0015 |

**Table 7.**
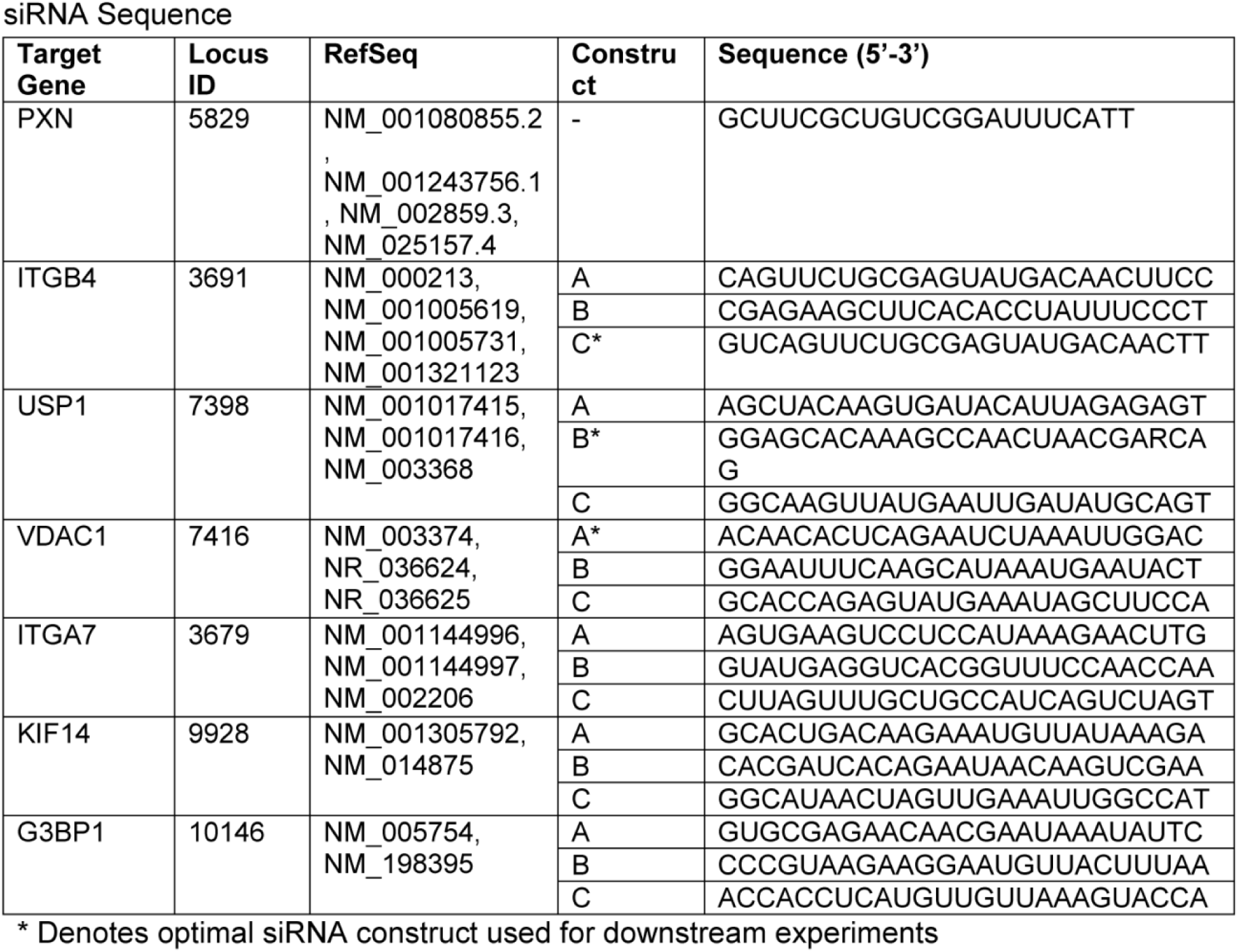
SiRNA sequences used for knockdown experiments

